# Phenotypic tolerance for rDNA copy number variation within the natural range of *C. elegans*

**DOI:** 10.1101/2025.03.21.644675

**Authors:** Ashley N. Hall, Elizabeth A. Morton, Rebecca Walters, Josh T. Cuperus, Christine Queitsch

## Abstract

The genes for ribosomal RNA (rRNA) are encoded by ribosomal DNA (rDNA), whose structure is notable for being present in arrays of tens to thousands of tandemly repeated copies in eukaryotic genomes. The exact number of rDNA copies per genome is highly variable within a species, with differences between individuals measuring in potentially hundreds of copies and megabases of DNA. The extent to which natural variation in rDNA copy number impacts whole-organism phenotypes such as fitness and lifespan is poorly understood, in part due to difficulties in manipulating such large and repetitive tracts of DNA even in model organisms. Here, we used the natural resource of copy number variation in *C. elegans* wild isolates to generate new tools and investigated the phenotypic consequences of this variation. Specifically, we generated a panel of recombinant inbred lines (RILs) using a laboratory strain derivative with ∼130 haploid rDNA copies and a wild isolate with ∼417 haploid rDNA copies, one of the highest validated *C. elegans* rDNA copy number arrays. We find that rDNA copy number is stable in the RILs, rejecting prior hypotheses that predicted copy number instability and copy number reversion. To isolate effects of rDNA copy number on phenotype, we produced a series of near isogenic lines (NILs) with rDNA copy numbers representing the high and low end of the rDNA copy number spectrum in *C. elegans* wild isolates. We find no correlation between rDNA copy number and phenotypes of rRNA abundance, competitive fitness, early life fertility, lifespan, or global transcriptome under standard laboratory conditions. These findings demonstrate a remarkable ability of *C. elegans* to tolerate substantial variation in a locus critical to fundamental cell function. Our study provides strain resources for future investigations into the boundaries of this tolerance.

## INTRODUCTION

Repetitive DNA contributes substantially to genomic variation, evidenced by the observations that approximately half of the human genome is repetitive and repetitive DNA tends to show higher mutation rates than nonrepetitive DNA (1–7). Nevertheless, the contribution of repetitive DNA to phenotypic variation has long been overshadowed by a focus on single nucleotide variation (7,8). One of the largest repetitive loci in the genome is the ribosomal DNA (rDNA), a genomic element of considerable biological consequence (9). rDNA encodes the ribosomal RNAs, the most abundant transcripts in eukaryotic cells; rRNAs form both essential structural components of the ribosome as well as the core of its catalytic function (10,11). rDNA also serves as the organizing center for formation of the nucleolus, an organelle with roles in the regulation of stress response, apoptosis, and the cell cycle (12–14). The rDNA transcriptional unit is encoded in arrays of tandemly repeated copies in nearly all eukaryotes (15). Here, we use “rDNA” to refer to the array encoding the largest pre-rRNA transcript, which will be processed into the 18S, 5.8S, and 26S/28S rRNAs.

The repetitive nature of the rDNA leaves it prone to copy number variation through unequal sister chromatid exchange, mitotic recombination, and DNA repair processes (16–20). Accordingly, vast natural variation in rDNA copy number has been observed in many species, including between individual humans and among strains of model organisms such as *Saccharomyces cerevisiae*, *Caenorhabditis elegans*, *Drosophila melanogaster*, and *Arabidopsis thaliana* (6,21–27). Conservative estimates report ranges of 100-600 copies per haploid genome for individual humans (27), 50-500 for yeast strains (28), and 70-400 copies per haploid genome for wild worm isolates (21,22). This variation can amount to differences of millions of base pairs between individuals. The importance of maintaining a certain copy number is supported by the stability of copy number in model organisms and reported phenomena of restoration of rDNA copy number after reduction (21,22,29–33).

The high rate of rRNA transcription in growing cells is feasible in part due to the multicopy nature of the rDNA (10,34), and yet the number of rDNA copies in many species appears to be in far excess of the amount required for ribosome biogenesis (34,35). Copy number variation outside of severe deletions does not usually correlate with rRNA expression (36–41). Severe reductions in rDNA copy number can restrict ribosome biogenesis, with detrimental effects on fitness, growth, and viability (42–46). The severity of phenotypic consequences tends to scale with the severity of rDNA copy number reduction (33,47). In *S. cerevisiae*, while reduction of rDNA copy number from the wild type ∼170 copies to ∼35 copies does not negatively affect indicators of ribosome biogenesis, it does alter genome replication timing and increase sensitivity to DNA mutagens (36,37).

Studies testing the phenotypic consequences of rDNA copy number variation within the natural range in an otherwise isogenic background are rare. *S. cerevisiae* offers one example with the observation that copy number reduction within the natural range of wild yeast alters Sir2-mediated silencing at the rDNA, telomeres, and mating-type loci (48). In humans, association studies have found that rDNA copy number variation among individuals correlates with differences in global transcriptome and mitochondrial DNA abundance (25). Recent studies report that human rDNA copy number variation also associates with blood composition, markers of kidney function, and adult body mass, but not adult tissue rRNA expression (49,50).

Understanding the relationship between natural-range rDNA copy number variation and phenotype becomes critically important in light of an increasing body of literature associating rDNA copy number changes or instabilities with various human pathologies. Many types of cancer have associations with rDNA copy number alteration or instability (51–56), and high rDNA copy number in smokers may indicate increased risk of lung cancer (57). rDNA instability and copy number differences have been reported with aging in yeast (58,59) as well as in some aging mammalian tissues (60–63). rDNA instability also associates with neurodegeneration (64). The nature of the relationship between inherent rDNA copy number and predisposition for certain traits has not been studied in detail. Exploring the contribution of rDNA copy number variation to phenotype in an otherwise isogenic model system could help further our understanding of the direct effects of this pool of existing genetic variation. The technical difficulty both of characterizing and manipulating this repetitive array has made such studies challenging (22,65).

*C. elegans* is a model system well-suited to exploration of the phenotypic impacts of rDNA copy number variation. Unlike many species, *C. elegans* hosts its 45S rDNA array in only one locus, promoting ease of genetic manipulation (66). Unlike *Drosophila* rDNA, the *C. elegans* rDNA is present on an autosome (chromosome I) next to the telomere and lacks the confounding factor of fly retrotransposon interruption (67,68). *C. elegans* furthermore is an androdioecious species, enabling both reproduction through self-fertilization and through male-hermaphrodite genetic crosses (69). Substantial variation in rDNA copy number exists among wild worm isolates, with the forty best-characterized wild isolates of *C. elegans* having anywhere from ∼70 to ∼420 copies of the 45S transcriptional unit (21,22).

Here, we leverage this natural variation and the genetic capabilities of *C. elegans* to generate resources and explore phenotypic relationships with rDNA copy number. We constructed a set of recombinant inbred lines (RILs) between two genotypes: the lab strain and a wild isolate with over three times as much rDNA as the lab strain. We find that rDNA copy number is surprisingly stable in these strains over 20 generations, demonstrating a lack of copy number reversion. We further introgressed wild isolate-derived rDNA arrays that span a ∼5-fold range in copy numbers into the lab strain background to generate a panel of near isogenic lines (NILs). With these we demonstrate that rDNA copy number does not correlate with steady-state rRNA levels. We observe no impact of rDNA copy number differences on lifespan, competitive fitness, or early life fertility. RNA-sequencing reveals minimal changes in the transcriptome between strains with different rDNA copy numbers. Together, these data suggest many life history traits in *C. elegans* are permissive of even drastic changes to rDNA copy number, including increases in copy number, indicating a substantial capacity to tolerate this degree of genetic disparity.

## RESULTS

### Recombinant inbred lines of *C. elegans* with two different rDNA copy number alleles do not demonstrate rDNA reversion

We aimed to create tools to understand genomic interaction with rDNA copy number and explore relationships between rDNA copy number and phenotype. To this end, we generated a panel of RILs using *C. elegans* wild isolate MY1, a strain with over 400 rDNA copies per chromosome I array (21,22,47). The 118 RILs in the final panel were selected such that approximately half of strains had each of the two parental rDNA copy numbers. These RILs present a resource for investigations into rDNA copy number contributions to specific traits, either additively or as an interacting locus. rDNA copy number of the MY1 wild isolate has been characterized in the range of 396-431 copies of the 45S rDNA locus, hereafter referred to as the 417-copy rDNA allele (21,47). MY1 also differs from the laboratory strain N2 by approximately 90,000 single nucleotide variants (SNVs) and 30,000 small insertion/deletions (indels) (21). None of these polymorphisms are present in the 45S rDNA sequence, however (21,45). N2 and its derivatives have on the order of 86-133 rDNA copies; the specific derivative strain used here, SEA51, will be referenced as the 130-copy allele (21,47). SEA51 was used because it carries a GFP-containing transgene (*mIs13*) on the right arm of chromosome I, proximal to the rDNA. *mIs13* served as a visual linked marker for the rDNA locus: the lower, 130-copy allele was linked to GFP presence and the higher, 417-copy allele, linked to GFP absence. To generate RILs, MY1 hermaphrodites were crossed with SEA51 males, GFP(+) F1s were selected and allowed to self-fertilize, and 118 F2s of either GFP(+) or GFP(-) phenotype were independently propagated to generate homozygous lines (**Figure 1A**, Methods).

**Figure 1:**
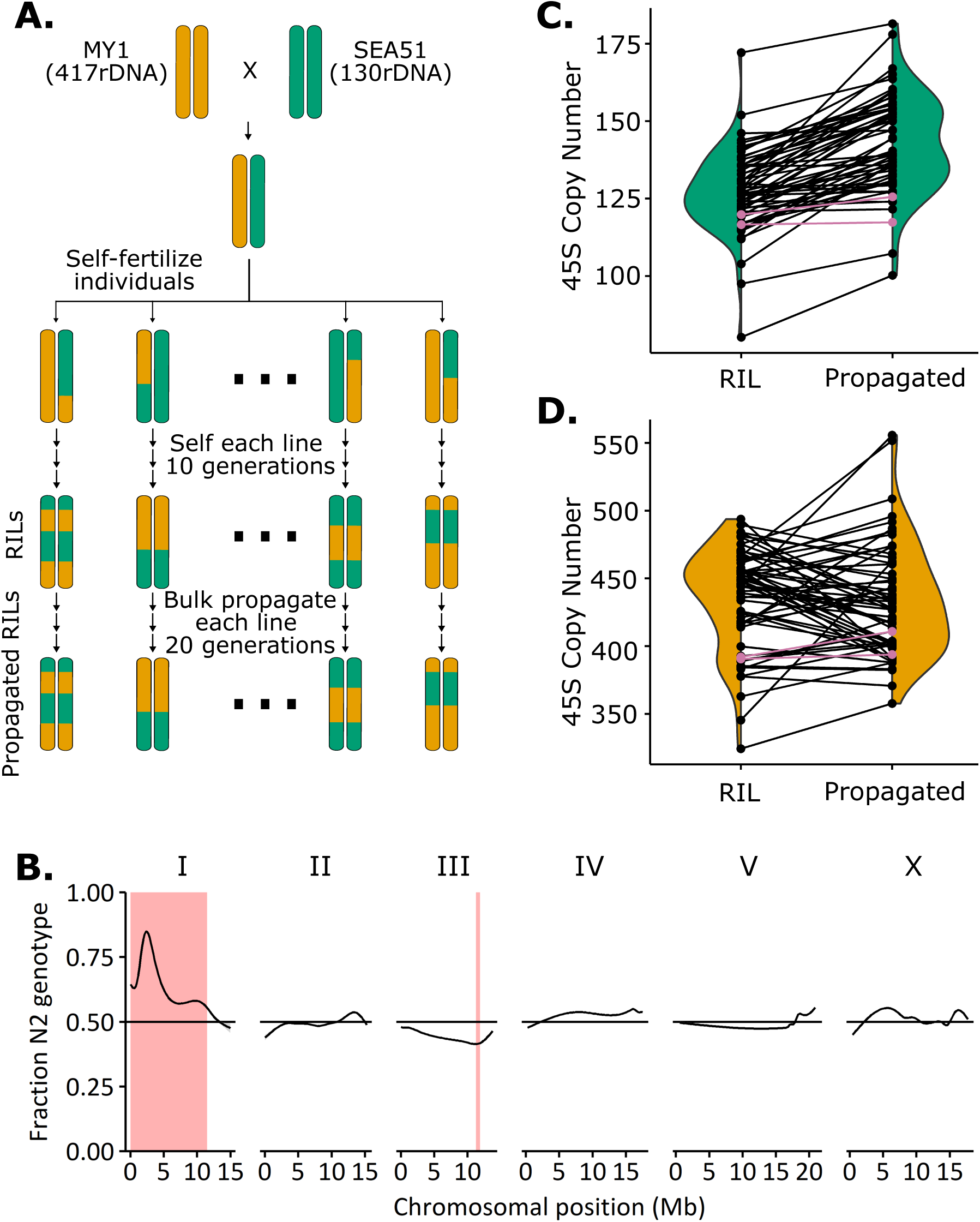
MY1 x SEA51 recombinant inbred line construction, genotyping, and rDNA copy number estimation. A: Parental strains MY1 (hermaphrodite) and SEA51 (male) were crossed and F1s were self-fertilized to initiate production of 118 RILs. At the F2 generation, 60 GFP(+) (130-rDNA) and 60 GFP(-) (417-rDNA) worms were selected to self-reproduce for 10 generations. After 10 generations of single-worm selfing, worm strains were frozen and designated as RILs. The RILs were then propagated an additional 20 generations in bulk. **B:** Proportion of RILs with N2 genotype at each position of the *C. elegans* genome. Regions highlighted in red are significantly distorted from a 50:50 ratio with a Bonferroni corrected p-value of 0.05 or lower. The peak on chromosome I at approximately 2.3 Mb matches the location of the *zeel-1*/*peel-1* toxin-antitoxin locus. **C:** rDNA copy number estimates from short read sequencing for RILs and propagated RILs linked to *mIs13*. The changes in copy number over the 20 generations of propagation were consistent with a normal distribution (Shapiro-Wilk, p=0.17) and no lines rose above the 200-copy number level (see **Figure S1B**). Pink lines indicate copy number estimates of the included SEA51 parental strain controls. **D:** rDNA copy number estimates from short read sequencing for RILs and propagated RILs not linked to *mIs13*. The changes in copy number over the 20 generations of propagation were consistent with a normal distribution (Shapiro-Wilk, p=0.08) and no lines fell below the 300-copy number level (see **Figure S1B**). Pink lines indicate copy number estimates of the included MY1 parental strain controls.

RILs were genotyped by short read sequencing; haploblock and rDNA copy number estimation were calculated from these data. Complete co-segregation of *mIs13* and the rDNA array was observed: every strain that was selected as homozygous GFP(+) had between 81-177 rDNA copies in the initial sequencing (**File S1**). Most chromosomes of a given RIL had between one and four large haplotype blocks, as expected from the cross strategy (**Figure S1A, File S2**). Observations of segregation distortion on chromosome I (**Figure 1B**) and at the mitochondrial DNA were consistent with predictable outcomes of known inheritance and wild isolate genetic incompatibility (70). All RIL mitochondrial genomes were MY1 genotype (**File S2**), as expected from the cross design (**Figure 1A**). An additional strong segregation distortion observed on chromosome I was likely the result of the well-documented *zeel-1*/*peel-1* toxin-antitoxin locus, present at 2.3Mb on chromosome I (70). MY1 is *zeel-1*/*peel-1* null and therefore incompatible with the SEA51 *zeel-1*/*peel-1* wild type locus (70), resulting in selection for SEA51 genotype on the left arm of chromosome I (**Figure 1B**). The right arm of chromosome I is not affected by this *zeel-1*/*peel-1*-driven segregation distortion due in part to the *mIs13*-based selection for equal representation of parental genotypes (**Figure 1B, S1A**).

Whole genome sequencing-based rDNA copy number estimations tend to have high error rates and batch-based variation (22). We took six RILs marked as having inherited the 130-rDNA array and validated their rDNA copy number using the lower-throughput but more reproducible method of contour-clamped homogeneous electric field (CHEF) gel followed by Southern blotting (**Figure S2A** and **Table S1**) (22). All six strains validated for inheritance of the 130 rDNA copy number range. Four of the six strains whose WGS estimates suggested differences of 7-26 copies among them yielded CHEF results that were all within a single copy of each other (118 or 119 copies), suggesting that some, but not all, of the variation in WGS copy number estimates may be artifactual and some copy numbers are even more uniform than apparent (**Figure 1C**).

The creation of these RILs allowed us to ask if extreme rDNA copy number is genetically dictated or if such arrays are maintained stably regardless of background genotype. We hypothesized that if the MY1 genetic background either requires or dictates large rDNA copy numbers, we would see some GFP(+) 130-rDNA lines substantially gain back rDNA copies and some GFP(-) 417-rDNA lines substantially lose copies over a propagation period. To allow opportunity for copy number mutation and selection to occur, we performed population propagation of the RILs for 20 generations and again used short-read WGS for rDNA copy number re-estimation (**Figure 1C, D**). We did not observe restoration of extremely high or low rDNA copy number among the propagated lines. The ranges of the two categories of RILs remained completely non-overlapping: the starting range of the homozygous GFP(-) lines was 324-494 copies and the post-propagation range was 358-556 copies. The starting range of the homozygous GFP(+) lines was 80-172 copies and the post-propagation range was 100-181 copies (**Figure 1C, 1D, S1B**). The distributions of copy number changes were consistent with a normal distribution for both sets of RILs (GFP(+) RILs, which inherited low copy number, and GFP(-) RILs, which inherited high copy number) (**Figure 1**), as would be expected for changes stemming from random fluctuation or measurement error. Most of the GFP(+) propagated RILs (48 out of 58) had rDNA copy number estimates increase by 5 copies or more over their unpropagated estimates, as did one of the SEA51 control strains, suggesting this increase may be in part systematic sequencing estimate error (see **Figure S1B**), although not ruling out the possibility of genuine copy number shift. However, none of the propagated RILs exhibited copy number changes that approached bridging the ∼3-fold distance between the parental copy numbers (**Figure S1B**). The absence of drastic rDNA reduction or amplification over the 20 generations of laboratory propagation suggests a degree of rDNA copy number stability that prompted us to attempt further rDNA copy number introgressions into the laboratory strain background.

### Near isogenic lines were created to investigate rDNA-specific influence on phenotype

In order to directly investigate potential phenotypic effects of rDNA copy number variation, we introgressed rDNA arrays from four wild isolates of *C. elegans* into the N2 derivative laboratory strain background (**Figure 2A**). Wild isolates were chosen from the highest and lowest ends of the spectrum of verified rDNA copy numbers – those that had been validated by two different estimation methods (WGS and CHEF gel) (**Figure S2**) (22). These wild isolates included JU775 and MY16 at the low end of rDNA copy number (**Figure S2B**); these alleles will be referred to as the 81 and 73 rDNA arrays, respectively (47). High rDNA copy number alleles were donated from wild isolates MY1 and RC301, both of which have over 400 rDNA copies (**Figure S2C**); these two alleles will be referred to as the 417 and 420 rDNA copy arrays, respectively. We generated near isogenic lines (NILs) using these four rDNA arrays (Methods). The 417-rDNA array was introgressed with further backcrossing of one of the above RIL lines. The other three rDNA arrays were introgressed in two iterations, first generating a set of NILs with >1Mb of wild isolate DNA remaining linked to the rDNA and then further refining the NILs through additional outcrosses and SNP-based genotype screening to select desired recombinants. **Figure 2** and **Table 1** detail the rDNA copy numbers and remaining linked wild isolate genotypes of each NIL (full data in **Files S3-S9**). rDNA copy numbers in resulting strains were verified by CHEF gel (**Figure S2**, **File S10**).

**Figure 2:**
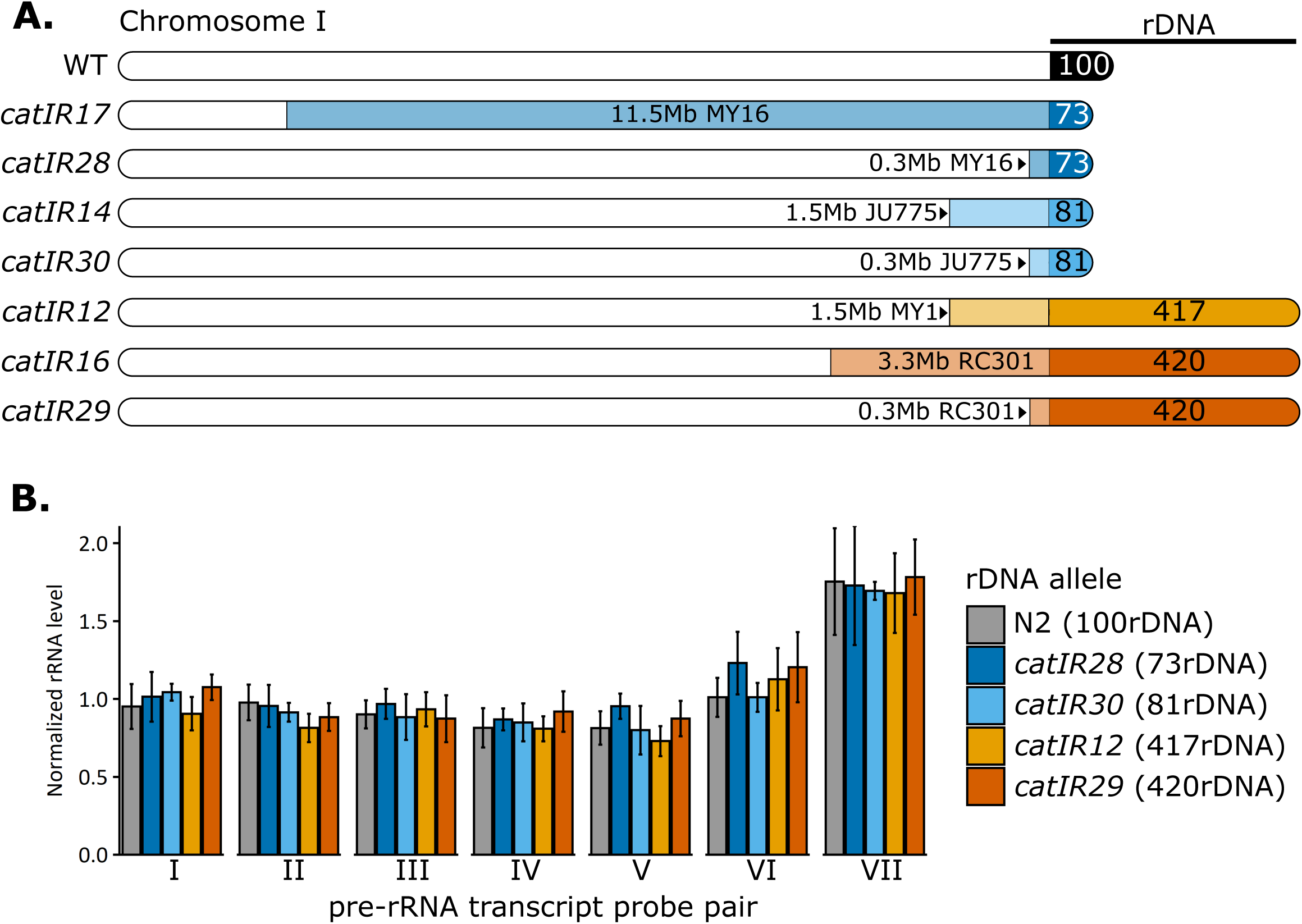
Near Isogenic Lines (NILs) with different rDNA copy numbers do not differ in rRNA expression. A: The 45S rDNA array in *C. elegans* is located on the far right arm of chromosome I. The schematic depicts the chromosome I genotype for each NIL generated. Allele designations for the introgressed region of each NIL are indicated to the left. The relative amount of wild isolate DNA remaining linked to the rDNA array is indicated by the lighter colored regions, next to the strain name of the source wild isolate (MY16, JU775, MY1, or RC301). The copy number of rDNA units is noted in the darker color regions for each NIL. Wild type (WT) is indicated for reference, with no linked wild isolate DNA and approximately 100-130 rDNA copies in N2 and its derivatives. White chromosomal regions correspond to N2 genotype. **B:** pre-rRNA levels from the 45S transcript were quantified by RT-qPCR in NILs with minimal wild isolate DNA. Seven different probe pairs were used across the transcript template (see **Figure S3G** and **Table S7**). rRNA levels were normalized to actin mRNA. No significant differences are present between any NILs and N2 (ANOVA with Tukey Honest Significant Difference test). See **Figure S3** for further rRNA quantification.

**Table 1:** Near Isogenic Lines of *C. elegans*.

| Strain | Genotype | rDNA copy number | Remaining linked SNVs* | rDNA designation |
| --- | --- | --- | --- | --- |
| SEA300 | *catIR12* [MY1 chrI:13527418-end] | 417 | 51 | 417-rDNA |
| SEA302 | *catIR14* [JU775 I:13529383-end] | 81 | 2,485 | 81-rDNA |
| SEA304 | *catIR16* [RC301 I:11737213-end] | 420 | 87 | 420-rDNA |
| SEA305 | *catIR17* [MY16 I:3576802-end] | 73 | 4,192 | 73-rDNA |
| SEA328 | *catIR28* I [MY16 I:14764185-end] | 73 | 12 | 73-rDNA |
| SEA329 | *catIR29* I [RC301 I:14989978-end] | 420 | 0 | 420-rDNA |
| SEA330 | *catIR30* I [JU775 I:14775350-end] | 81 | 220 | 81-rDNA |

\*Number of variants from GATK pipeline, ignoring blacklisted regions. While this shows no variants in SEA329, RC301 genotype was confirmed at position 14989978 as part of the strain construction process. For MY1 NILs, listed is the number of N2 variants linked to the rDNA. Values are approximate.

### rRNA levels do not reflect rDNA copy number

A straightforward mechanism by which rDNA copy number variation could impact phenotype is through altered rRNA abundance. However, reports from several species suggest that once past a minimal threshold, differences in rDNA copy number do not correspond to differences in rRNA levels (36,39,71,72). We measured rRNA levels in synchronized Day 1 adult worms of our NILs containing introgressed rDNA copy number variants. Steady-state levels of 18S and 28S mature rRNAs did not differ in an rDNA copy number-dependent manner (**Figure S3A-E, File S11-S12**). While the 5S rRNA is encoded in a separate array on a different chromosome from the 45S locus, 5S transcripts must still be incorporated into the ribosome in equimolar amounts to the 45S products; we therefore additionally assessed 5S rRNA. There were no differences in 5S rRNA levels among the strains tested (**Figure S3F, File S13**). Transcripts from the 45S pre-rRNA also did not differ in a copy number-dependent manner, with no pre-rRNA regions differing significantly between any NIL and wild type (**Figure 2B**, **Figure S3G, H, File S13**). Overall, these data support the notion that rRNA transcription is not regulated by rDNA copy number variation in *C. elegans* within the range investigated, but rather by other regulatory mechanisms (73).

### rDNA copy number does not correlate with competitive fitness

Competitive fitness assays represent an integration of many aspects of life history (growth, development, fertility) and allow for small fitness defects to compound over generations. We performed pairwise competitive fitness tests of a GFP-marked laboratory strain (SEA51) and our highest copy number NILs, which represent the greatest divergence from the control strain copy number (a >3-fold difference in copy number). The *mIs13* transgene was used in the competitions to trace worms with the wild-type rDNA allele (∼130 copies). Expression of this transgene alone causes a fitness defect: the SEA51 strain, when competed against wild type N2, decreased to 15% of the population after propagation for ∼10-11 generations (**Figure 3, Figure S4A-C, File S14**). To account for this defect, we assessed relative competitive fitness in NIL competitions compared to control competitions. To confirm that final population proportions did not arise from a chance event early in the experiment, we also verified approximately equal population proportions at an early timepoint (**Figure S5**).

**Figure 3:**
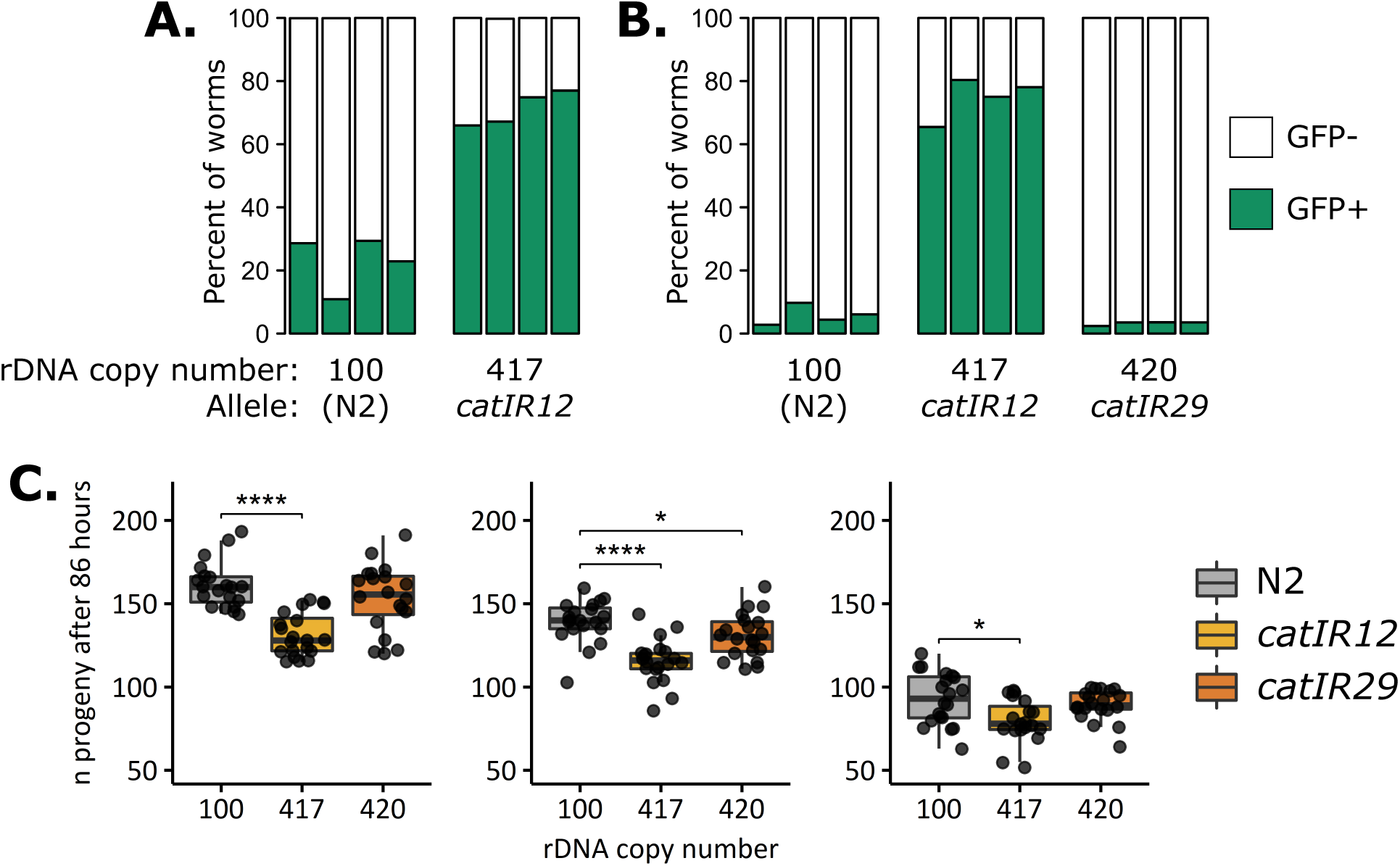
rDNA copy number variation does not correlate with reduced competitive fitness or early life fertility. A-B: Competitions were conducted between strains with high rDNA copy number (GFP(-); >400 rDNA copies, alleles indicated at bottom) and marker strain SEA51 (GFP(+); 130 rDNA copies). **A:** Four independent replicates were conducted each of either competition between N2 and SEA51 or between the 417-rDNA NIL (allele *catIR12*) and SEA51, experiments propagated simultaneously. **B:** Four replicates each were conducted of competitions between SEA51 and either N2, the 417-rDNA NIL (allele *catIR12*), or the 420-rDNA NIL (allele *catIR29*), experiments propagated simultaneously. For all panels, bars represent the relative proportion of worms that are GFP(+) (green, SEA51) or GFP(-) (white) after ∼10-11 generations. At least 1,000 worms were quantified for each bar (**File S14**). **C:** Early life fertility was assessed for N2 and NILs with high rDNA copy number in three replicates, n=20 individual worms per strain per replicate. Data were collected from 86 hours post-embryo of the worm’s life and thus represent progeny production through just the first day of reproductive adulthood. The data fail the Shapiro-Wilk normality test and are not normally distributed. Statistical tests represented in the figure are Pairwise Wilcoxon tests with Benjamini-Hochberg were performed to compare strains. * p<0.1, **** p< 0.001.

Increasing rDNA copy number to >400 copies did not confer a consistent fitness defect attributable to rDNA; rather, a strain-specific fitness defect was observed. The 417-rDNA allele derived from MY1 (*catIR12*) had a severe fitness defect: SEA51 reproducibly outcompeted the strain with this allele (417-rDNA GFP(-), **Figure 3A, B, Figure S4C**), indicating an associated fitness defect more severe than that of the *mIs13* transgene. However, other strains with 420 rDNA copies did not exhibit this severe fitness defect. Two different 420-rDNA copy alleles from wild isolate RC301 competed either near-equally with the GFP control strain or outcompeted it to the same degree as N2 (**Figure 3B, Figure S4A, B**), implying that high rDNA copy number was not the causative factor. The strains with rDNA from RC301 differ in the amount of linked wild isolate DNA; the strain with less linked wild isolate DNA competed the same as N2 (**Figure 3B**). This suggests that the reduced competitive fitness comes from a non-rDNA SNV or indel native to the wild isolate backgrounds. Of all the strains with high rDNA copy number, the 420-rDNA *catIR29* allele has the fewest genetic variants linked to the rDNA (**Figures 2A**, **Table 1**). Taken together, these data suggest that high rDNA copy number did not causally reduce competitive fitness but rather linked non-rDNA variants reduced fitness to different degrees in the different strain backgrounds.

Because the competition assays were restricted to pairwise comparisons, we also directly quantified early life fertility in all of the NILs. We defined early life fertility as viable progeny produced within the first ∼24-30 hours of egg laying. The 417-rDNA NIL (allele *catIR12*) shows a consistent reduction in early life fertility (**Figure 3C, File S15**). We again do not attribute this fertility defect to high rDNA copy number, as it was not observed in the 420-rDNA *catIR29* high rDNA copy number strain (**Figure 3C**). Strains with reduced rDNA copy number also displayed no consistent early life fertility phenotype attributable to the rDNA (**Figure S6A**). Alleles with high amounts of linked wild isolate DNA for the same rDNA copy numbers did display some early life fertility defects (**Figure S6B**). The version of the 420-rDNA and 81-rDNA arrays that have high linked wild isolate DNA (*catIR16* and *catIR14*) exhibited reduced early progeny production compared to N2 (**Figure S6B**) that was not present for their outcrossed counterparts, consistent with the competition data and again suggesting causative linked variation that was resolved by the additional outcrossing. In summary, we conclude that rDNA copy-number variation did not causally affect early life fertility or competitive fitness.

### Changes in rDNA copy number did not alter lifespan

Lifespan is a highly multigenic trait. Changes in rDNA copy number, rDNA instability, and changes in nucleolar morphology have been linked to aging (63,74–77), leading us to question if altered genomic rDNA copy number would impact lifespan of worms. While our NILs did not differ in rRNA expression, there are other mechanisms by which differences in rDNA copy number could influence lifespan (36,37). We measured lifespan in the NILs and N2 and observed no impact of the rDNA copy number changes on lifespan. In only one replicate of one strain (73-rDNA, *catIR28*), lifespan was significantly shortened; in the other two comparisons of the 73-rDNA array and in all assays using copy numbers 420, 417, and 81, lifespan of each NIL did not differ from wild type (**Figure 4, Figure S7, File S16**).

**Figure 4:**
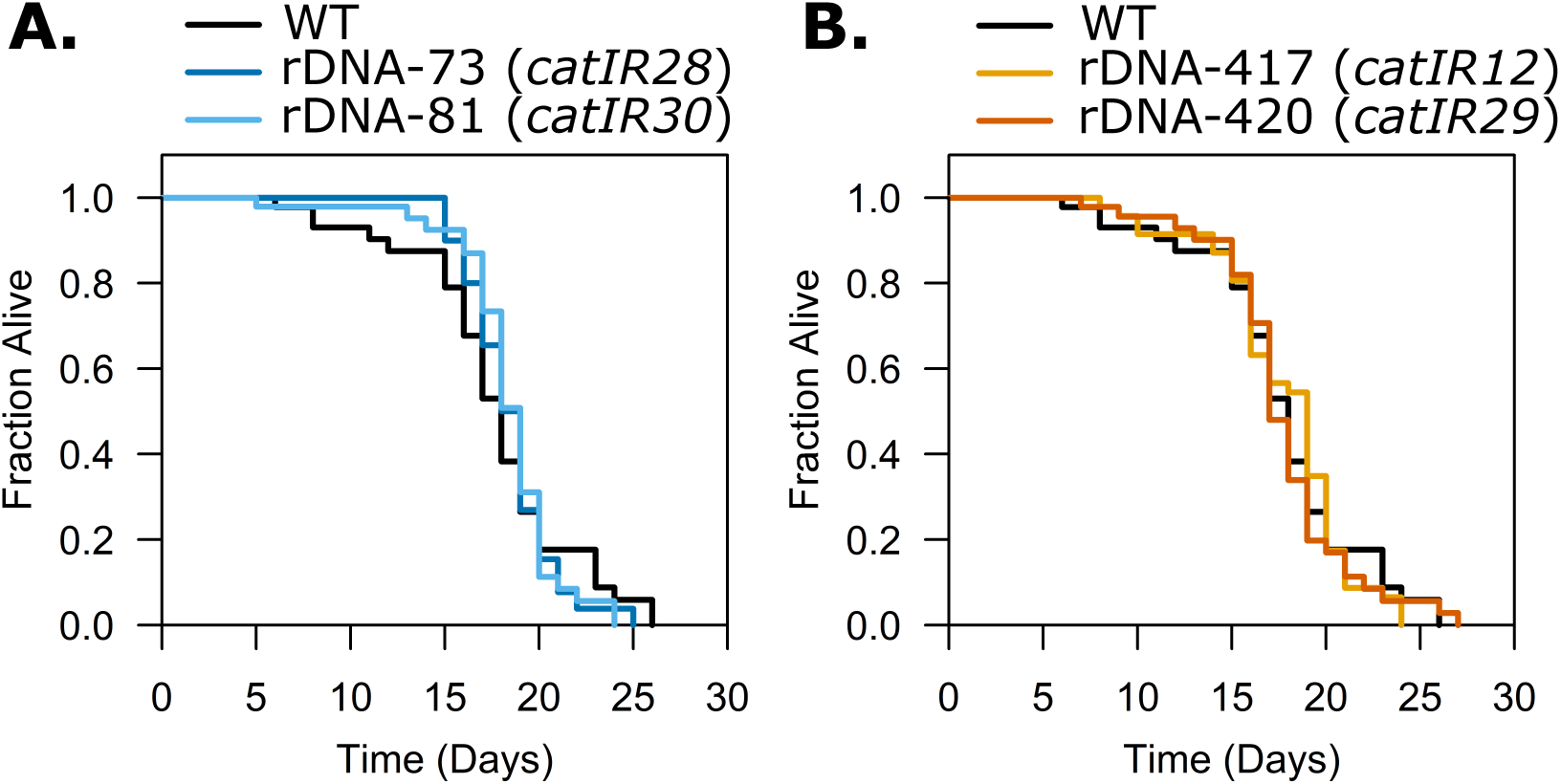
Lifespan of *C. elegans* is robust to increases or decreases in rDNA copy number within the natural range. Lifespan assays were conducted on N2 and NILs with various rDNA copy numbers. **A:** NILs with low rDNA copy number. **B:** NILs with high rDNA copy number. (n = 50)(**File S16**) Lifespans for all five strains in **A** and **B** were performed simultaneously (N2 data are the same between the plots).

### Changes in rDNA copy number did not alter the frequency of males in a sensitized background

Having sufficient rDNA copies is important for maintaining proper genome segregation. In *S. cerevisiae*, transcription of the rDNA controls levels of condensin binding such that more condensin binds to the rDNA when fewer repeats are transcribed (78), and transcription from the rDNA impacts chromosome segregation (79,80). Reduced rDNA copy number in yeast can also lead to premature cell cycle anaphase entry (36). We looked for signs of global nondisjunction in *C. elegans* NILs with rDNA copy number changes. To test for enhanced chromosome nondisjunction, the frequency of male progeny was assessed. Males arise through nondisjunction of the X chromosome (XO males vs. XX hermaphrodites) and in wild type worms, the frequency of male progeny is approximately 1 in 1000 (81). We used a sensitized strain background that has a high incidence of nondisjunction of both the X chromosome and of autosomes (*him-5*(*ok1896*)) (82). *him-5* is involved in meiotic break positioning and is a paralog of *rec-1* (83). The null allele *him-5*(*ok1896*) was introduced into the NIL backgrounds and strains were measured for the incidence of each type of nondisjunction progeny produced by this mutation: dead embryos (presumed autosomal nondisjunction), male worms (OX), dumpy worms (presumed XXX), and those characterized as having “other defects” (typically slow growth). There were no replicable, statistically significant differences between any NILs and wild type carrying *him-5*(*ok1896*) (**Figure S8, File S17**). Therefore, neither increasing nor decreasing rDNA copy number changes the frequency of meiotic nondisjunction events in the *him-5*(*ok1896*) mutant background.

### Global transcriptomes are highly similar between worms with different rDNA copy numbers

Global transcriptomic changes have been reported in flies with rDNA copy deletions (84), and *Arabidopsis thaliana* lines with reduced rDNA copy number also show changes in gene expression, including effects on metabolism and ribosome biogenesis (85). Neither the impact of increasing rDNA copy number nor the impact of altering rDNA copy number in near isogenic backgrounds of *C. elegans* have been explored. We performed RNA-sequencing on synchronized Day 1 adult worms of NILs containing minimal linked wild isolate DNA (**Figure 2A**). Two high rDNA copy number alleles, two low rDNA copy number alleles, and N2 were included in the assays. PCA revealed that all worm samples had similar transcriptomes (**Figure 5A**). Samples with the same rDNA copy numbers did not always cluster with each other, and few genes were differentially expressed when comparing any individual NIL to wild type (**Figure 5, Files S18-S21**). Gene ontology analysis of those genes that were differentially expressed in each NIL compared to wild type did not identify categories directly associated with rRNA expression or processing (**File S21**).

**Figure 5:**
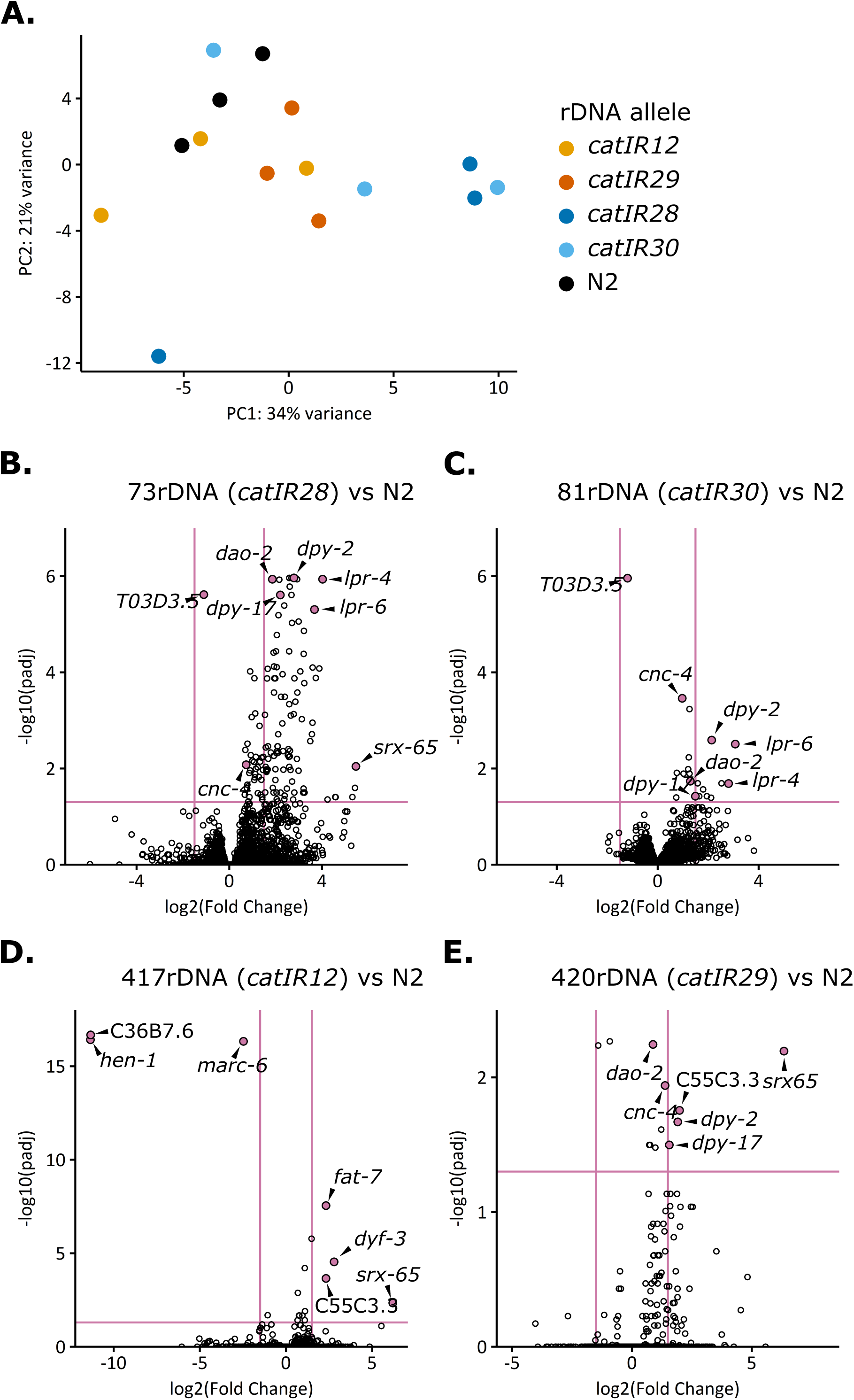
Few genes are expressed differentially between individual NILs and N2. A: Principal component analysis (PCA) was conducted on RNA-seq datasets from synchronized Day 1 adults of four NILs and N2. The rlog transformed DEseq dataset was analyzed by PCA in the DEseq package. Datapoints are colored according to the rDNA copy number allele. **B-E:** Three biological replicates for each NIL and N2 were used to calculate differential expression of all genes using DESeq2. Horizontal pink line indicates an adjusted p-value cutoff of 0.05, vertical pink lines indicate log2 fold changes of 1.5. Genes whose expression differences are shared among strains or have particularly high significance of large log2 Fold Change are indicated with gene name.

The strain with the most genes differentially expressed as compared to wild type was the NIL with 73 rDNA copies (*catIR28*) (134 genes, **Figure 5B**). Gene ontology analysis on the list of genes differentially expressed in the 73-rDNA strain as compared to wild type (**Tables S2-S4**) found enrichment of genes involved in cuticle biosynthesis, molting, and the extracellular region (note that while 13 cuticle constituent/development genes were differentially expressed in the 73-rDNA strain, four of those genes were also observed in the 81-rDNA strain dataset, and two in the 420-rDNA strain dataset, **File S21**). In agreement with the gene ontology analysis, genes associated with paralyzed, dumpy, molting, and movement variant phenotypes were enriched (**Table S3**), as were genes known to be involved in the epithelial system (**Table S4**). We performed a phenotypic assay of cuticle integrity in adult worms of N2 and the 73-rDNA NIL and found no difference in cuticle permeability between these two lines (**Figure S9, File S22**). An alternative hypothesis not tested here is that a difference in developmental timing of the *in utero* embryos of the 73-rDNA NIL worms accounts for the expression differences, rather than expression changes in the adult soma, consistent with the fact that the identified genes are known to be expressed in embryos. While the gene expression changes could conceivably be related to rDNA, the lack of congruence between the two lowest copy strains (73-rDNA and 81-rDNA) argues for allele-linked variation underlying the expression differences. Overall, we find that under standard laboratory conditions, rDNA copy number variation does not globally affect gene expression in Day 1 adult worms.

## DISCUSSION

In *C. elegans*, a single copy of the 45S rDNA unit measures 7.2kb; the difference in rDNA copy number between the highest and lowest ends of the natural spectrum, ∼70 to ∼420 rDNA copies, therefore equates to a difference of 2.5Mb of DNA per haploid genome – a considerable amount of DNA for an animal with a haploid genome of only 100 Mb (86). This DNA needs to be both replicated and regulated, introducing a discrepancy in genome maintenance load between strains. Surprisingly, this inequitable burden among strains did not impact the life processes we investigated here, pointing to a remarkable tolerance for this additional genomic content.

We observe that under standard laboratory growth conditions, rDNA copy number within the natural range does not associate with steady-state rRNA levels, competitive fitness, early life fertility, or lifespan in *C. elegan*s. In contrast to association studies from other organisms, we show negligible effects of rDNA copy number on the global transcriptome (25,84). We furthermore observed no extreme rDNA copy number shifts over the course of 20 generations of bulk propagation in our recombinant inbred lines. It is of course possible that our 20-generation propagation period was insufficient for rDNA copy number to both mutate and undergo a detectable degree of selection, so the full extent of rDNA stability and its conditions of alteration in worms remain to be determined. A recent study of rDNA copy number mutation in yeast suggested a rate of one copy lost per 67-120 generations (87). If rDNA behaves similarly during *C. elegans* cell divisions and an estimated ∼15 cell divisions occur between generations of *C. elegans* (88,89), a predicted 2-4 copies would be lost over 20 worm generations, well below what our methods have the ability to reliably detect (22). Nevertheless, relative stability of all 118 lines over this period of time shows that rDNA copy number is neither directly controlled by non-rDNA loci, nor subject to frequent large mutation or strong selection. Importantly, this degree of stability allowed introgression of wild-sourced rDNA arrays into near isogenic lines without major alteration of copy number (**Figure S2**)(47). Stability of rDNA copy number is consistent with some reports in flies and yeast of rDNA deletions failing to regain their original copy number even over extensive propagation (46,90). It is in contrast, however, to other reports of rDNA copy number maintenance or rapid copy number recovery after loss (30,31,33,91), although these losses were below the natural copy number range and subsequent gains restored natural-range copy numbers.

rDNA copy number did not correlate with rRNA expression in our NILs, an observation that was not unexpected, given observations in other species (36,39,71,72). The regulation of rRNA expression is complex, including transcriptional silencing by histone modification (92–94), DNA methylation (95), antisense ribosomal siRNAs (96), and regulation by RNA polymerase I transcriptional machinery (97,98), among other factors (73). Previous studies, largely in yeast, have established that the minimal requirements of rDNA copy number for ribosome biogenesis are well below the boundaries of the known natural range and drastic reduction must occur before ribosome function is impaired (34,36,37,47). Given the substantial difference in rDNA dosage in the NILs we created here, these strains may provide useful future tools for an exploration of the robustness, scope, and cost of this regulation. The characterization of the ribosome biogenesis and translational capacity of these lines under conditions of stress may offer further insight into the degree to which rDNA copy number informs these cell processes and sets cell limits. Environmental or genetic perturbations that introduce stress or dysregulation of any of these factors in our NILs could potentially expose consequences to rDNA copy number variation previously unapparent, a subject for future investigations.

An intriguing link between lifespan and nucleolar size has previously been established in *C. elegans* (74,99,100). Reduced nucleolar size is observed in long-lived mutants (74) and during adult reproductive diapause (99), and smaller nucleoli are predictive of increased lifespan in wild-type worms (74). Nucleolar size tends to correlate with increased rRNA production (13,74,99,101) but not in all circumstances (72,99). We did not quantify the nucleolar size of the NILs generated here; their unaltered rRNA load and lifespans would *a priori* predict unaltered nucleolar size. However, testing this hypothesis could provide valuable insight into the compensatory mechanisms for rDNA copy number variation and its role in phenotype.

Although there was reason to hypothesize that the burden of far too many or too few rDNA copies would influence the phenotypes we have investigated here, there are many more phenotypes not tested, including responses to different environmental conditions (33) or phenotypic models of disease or dysregulation. For example, further exploration of cuticle morphology and molting defects may be warranted (102,103). Given the emerging link between rDNA copy number and cancer, a test of tumor proliferation (104) in these lines may provide insight into rDNA copy number as a cancer risk factor.

Although many other *C. elegans* RIL and RIAIL (recombinant inbred advanced intercross line) panels have been created, to our knowledge ours is the first biparental cross using source strains with rDNA arrays validated at such vastly different sizes. The resources we develop here thus enable future studies into communication between variation across the genome and the rDNA array in different conditions. Due to the sequence divergence of MY1 and N2, the RIL panel can also be used for standard quantitative trait locus discovery of phenotypes associated with wild isolate MY1 (105). Recombinant lines using wild isolates (commonly Hawaiian isolate CB4856) have been useful for insight into genetic and phenotypic variation and evolution of *C. elegans* (106–111); MY1 has been characterized for its phylogenetic relationship to dozens of other wild isolates (112) as well as included in a 16-wild isolate multiparental experimental evolution panel (111). The RIL panel here presents the opportunity to study this strain’s suite of genetic variation specifically in two different rDNA copy number contexts (high vs. low), facilitating future studies into the potential for rDNA context-dependence in the effects of genetic variation under non-standard growth conditions or with respect to phenotypes not explored here.

Some traits showed consistent differences between NIL 417-rDNA (allele *catIR12*, sourced from MY1) and N2 (**Figure S6**), but we do not attribute these differences to the abundance of rDNA, due to the uniqueness of the defects to the 417-rDNA array and absence from the 420-rDNA array. Rather, these phenotypes are more likely due to non-rDNA variation specific to that strain. There are four unique missense variants in the 1.5 Mb region proximal to the rDNA native to the MY1 (417-rDNA) wild isolate parental genotype but not the RC301 (420-rDNA) genotype (**Table 2**). Of particular note may be the missense mutation found in *ekl-4* in MY1: this gene has phenotypes of slow growth and embryonic lethality when knocked down with RNAi (113), making it both a candidate for further investigation and potentially a naturally-occurring variant that nevertheless may be conferring disadvantageous phenotypes in the N2 background. These data alone provide a starting point for avenues of discovery into genetic variation and consequences of breaking epistatic relationships in wild isolates of *C. elegans*.

**Table 2:** Variants with predicted high impact present in MY1 and RC301 in the ∼1.5 Mb proximal to the rDNA.

| Gene | Amino Acid | Base pair* | Wild Isolate |
| --- | --- | --- | --- |
| W04A4.6 | 79V>79A | 13687336A>G | RC301 |
| Y105E8A.32 | 50R>50L | 14378647C>A | MY1 |
| *ekl-4* | 233R>233K | 14470756G>A | MY1 |
| Y105E8A.20 | 305S>305L | 14488002G>A | MY1 |
| Y105E8A.20 | 423S>423L | 14488002G>A | MY1 |
| *tag-4* | 213R>213W | 14989978A>T | MY1, RC301 |
| F31C3.3 | 377R>377Q | 15049340C>T | MY1 |

\*Base pair positions correspond to the WS276 genome and variants and their effects were identified with the CeNDR variant effect browser (114).

Overall, we have observed that complex phenotypes in *C. elegans* including fitness and lifespan are forgiving of upwards of a ∼5-fold difference in rDNA copy number. Although interindividual differences in lifespan occur between nominally isogenic worms (115), potentially through variation in translational capacity (116), our data argue that these differences are not attributable to the mutability of the rDNA array or individual variation in rDNA copy number. Our data further demonstrate that on a population level, rDNA copy number does not rapidly revert to a genetic background-prescribed copy number. The possibility remains that specific conditions will reveal phenotypes associated with rDNA copy number variants, but our results show that for the phenotypes assessed, worms are able to correct for and normalize phenotype across large differences in genomic content. These results speak to the magnitude of regulatory capacity available to the genome and paves the way for investigation into the mechanisms and timing of such accommodation.

## METHODS

### Worm husbandry and strain construction

*C. elegans* were grown at 20°C on NGM seeded with OP50 bacteria unless otherwise specified. Male stocks of worms for crosses were made by incubating L4 hermaphrodite worms at 30°C for four hours, then returning plates to 20°C and screening for male progeny 3-4 days later. Crosses were conducted by combining 10 male worms with three L4 hermaphrodite worms on 3cm NGM plates. Successful mating was ascertained by the presence of approximately 50% male progeny in the F1 generation. Some crosses utilized transgene *mIs13* as a GFP marker, in these cases, F1 cross progeny were selected based on the presence or absence of pharyngeal GFP. *mIs13[myo-2p::GFP + pes-10p::GFP + F22B7.9p::GFP]* is approximately ∼615kb in size encoding ∼102 copies of GFP, expressed constitutively in the pharynx, intestine, and germline (117). Strain SEA51 was generated by outcrossing PD4788 to N2 (VC2010) six times, followed by three generations of single-worm propagation. Strains used are listed in **Table S5**.

### NILs

Initial stages of NIL generation used the *mIs13* GFP marker to track rDNA alleles. Once past the stages at which GFP marker could be used, an rDNA-proximal restriction fragment length polymorphism or insertion/deletion that differed between N2 and the wild isolate of interest was used as a genotyping marker to follow rDNA copy number alleles (**Table S6**). Construction of SEA300 (417-rDNA) was performed by crossing SEA51 males to hermaphrodites of RIL SEA159 (GFP(-), >400 rDNA); the backcross to SEA51 males was performed sequentially a total of six times, followed by six generations of individual self-reproduction to create an initial introgressed line. Males were produced from this line and crossed to SEA51 GFP(+) hermaphrodites; GFP(-) F2s were selected from this cross to generate strain SEA300, containing 417-rDNA and wild-type mitochondrial DNA. Construction of NILs SEA302, SEA304, and SEA305 was performed using wild isolates, JU775 (81-rDNA), RC301 (420-rDNA), and MY16 (73-rDNA), respectively. Male SEA51 GFP(+) worms were crossed to wild isolate hermaphrodites and GFP(+) F1 progeny were allowed to self-fertilize. GFP(-) F2 progeny were crossed to SEA51 males again, for a total of 8 SEA51 crosses. The last backcross was to a SEA51 hermaphrodite to restore wild-type mitochondrial DNA. After the final cross, progeny were propagated as GFP(-) single worms for six generations. The resulting NILs are diagramed in **Figure 2A** and exhibit >1Mb of linked wild isolate DNA. To generate NILs with less linked wild isolate DNA, SEA302, SEA304, and SEA305 were further outcrossed to N2 two to three more times with genotype screening using RFLP or indel markers to produce NILs SEA330 (81-rDNA), SEA329 (420-rDNA), and SEA328 (73-rDNA) (**Table 1**).

### Worm synchronization by hypochlorite treatment (bleaching)

Gravid adult worms were washed from plates into 15mL conical tubes in 1X M9 [42mM Na_2_HPO_4_, 22mM KH_2_PO_4_, 22mM NaCl, 1mM MgSO_4_] and pelleted either by gravity settling or by centrifugation, followed by removal of the supernatant. Worms were lysed with ∼8-10mL of a hypochlorite solution (0.5N NaOH, ∼0.8%-1.2% NaOCl) with agitation for no more than seven minutes followed by embryo collection by centrifugation (1500 rpm for 1 minute, with slow deceleration). Embryos were washed 3x in 10-15mL 1X M9. Embryos were then either plated on NGM+OP50 or, for stage synchronization, on unseeded NGM plates with a layer of ∼5mL 1X M9 covering the plate. For stage synchronization, embryos were allowed to hatch in the 1X M9 overnight and starve as L1 larvae. Larvae were harvested by centrifugation in 15mL conical tubes (1500 rpm for 1 minute) and washed once in 1X M9 before being plated on food.

### RIL construction and propagation

Male SEA51 worms were crossed to hermaphrodite MY1 worms and cross progeny were selected by presence of GFP (transgene *mIs13[myo-2p::GFP + pes-10p::GFP + F22B7.9p::GFP]*). Eighty GFP(+) F1s were allowed to self-fertilize on plates in populations. 120 GFP(+) (130-rDNA) and 72 GFP(-) (417-rDNA) F2 worms were picked to single wells of 24-well NGM plates. F3 worms were screened to confirm homozygosity of the F2 GFP(+) worms. 60 GFP(+) and 60 GFP(-) individual homozygous F3 worms from separate wells were propagated to new wells of 24-well plates. These 120 worm lines were propagated as single worms every generation for 10 generations. In each propagation, two individual worms were picked to separate wells for each line; only one of the two wells would be used in the next generation of propagation, while the other well was a backup against loss of individual worms (such as from crawling up the walls of the plate). Even with these measures, two worm lines were lost during propagation, resulting in 118 recombinant inbred lines (RILs) at the end of propagation. At the F12 generation of worms (10 generations of single-worm descent) worm populations were grown in bulk for each line and both frozen and collected for whole-genome sequencing. RILs were then propagated an additional 20 generations in bulk. For bulk propagation, worms were grown to gravid adulthood until many embryos were laid. Adult worms were washed off of the plates with 1X M9, leaving behind the embryos. A 1cm x 1cm chunk of agar (at least 100 embryos) was cut out and propagated to a new 6cm NGM plate. This process was repeated every three days for 20 generations. After 20 generations, worms were propagated to 10cm NGM plates and allowed to starve for both strain preservation (freezing) and harvest for genomic DNA preparation.

### Genomic DNA Isolation

A Qiagen DNeasy Blood & Tissue DNA purification kit (69504) was used for genomic DNA extraction. Worm pellets frozen in kit-provided ATL buffer were freeze-thawed three times between −20°C and 37°C. 20μL proteinase K was added and the samples were incubated at 56°C for 3hr with occasional vortexing. 4μL 100mg/mL RNase A (Qiagen 19101) was added to each sample and incubated at room temperature for 5 minutes. 200μL AL buffer was added, and the DNA extraction continued as described in the kit protocol. Final DNA was eluted in a total volume of 100μL. DNA concentration was determined by Qubit assay for sequencing library preparation.

### Short-read sequencing library preparation and rDNA copy number estimation

Libraries were prepared largely as published using 10ng DNA as input (22). Briefly, the Illumina Nextera DNA Sample Preparation kit (FC-121-1030; discontinued) was used. KAPA HiFi 2x Master Mix was substituted for Illumina NPM master mix in the PCR amplification steps. The 10ng DNA was tagmented in a 20μl reaction containing 1μl tagmentation enzyme and 10μl tagmentation buffer and incubated at 55°C for 8 minutes. The reaction was stopped by addition of 10μl 5M guanidine thiocyanate and DNA was purified with AMPure XP beads (Beckman Coulter A63881) with the following ratio: 15μL AMPure beads and 25μL binding buffer [20% PEG8000, 2.5M NaCl] to the now 30μL reaction. Dual-barcoded libraries were PCR amplified in a 22.5μL reaction containing 10μL tagmented DNA, 7.5μL Kapa HiFi 2X master mix and 2.5μL of each Illumina barcode indexing primer. PCR conditions were as follows: 72°C 3 min, 98°C 30 sec, 98°C 10 sec, 63°C 30 sec, 72°C 40 sec, with the latter three steps cycled 6 times. The post-PCR libraries were purified with 30μL AMPure XP beads. Final libraries were quantified by Qubit high sensitivity assessment (Invitrogen Q32854) and diluted to 2nM. Libraries were denatured and diluted for sequencing as per the manufacturer’s instructions (NextSeq Denature and Dilute Libraries Guide 15048776 Rev. D) and libraries were sequenced with a 75bp-paired end NextSeq 500/550 High Output v2 150 Cycle kits (FC-404-2002).

For RILs, mixed-stage starved larvae were used for gDNA preparation and sequencing libraries were prepared in two batches for each the RILs and propagated RILs. With each library preparation batch, a control sample of SEA51 DNA and MY1 DNA was prepared in the same batch. Short-read genome sequencing of RILs was conducted at three points: as soon after initial F2 selection as a population could be grown up (collected worms approximately F4 animals, Phase 1), following the completion of single-worm propagation (collected worms approximately F14, Phase 2), and following completion of 20 generation of bulk propagation (collected worms approximately F33, Phase 3) (**File S1**). rDNA copy number was estimated using a maximum likelihood estimation method (**File S1**) (22,118,119). Genotypes were determined with GATK HaplotypeCaller and the map was filled with the max marginal method (**File S2**). For N2 NILs, libraries of mixed stage worms were prepared for genotyping of each NIL. VCF files of chromosome I of NILs available in **Files S3-S9**.

### CHEF plug sample preparation, run conditions, and Southern blotting

CHEF plugs were prepared in a similar manner to previous studies (22), with the following changes: worms were grown up either by placing ten L4 or Day 1 adult worms on a 10cm NGM + OP50 plate and growing to starvation (∼5-6 days at 20°C); for select assays a single L4 or Day 1 adult worm was placed on a 6cm NGM+OP50 plate and produce a population to grow to starvation (∼5-6 days at 20°C). Starved worms were washed from plates into 15mL conical tubes or 1.5mL microcentrifuge tubes, pelleted, and washed in 1X M9 buffer. Worms were washed one time in autoclaved double glass distilled water, and the volume of worms and water after removing the wash supernatant was estimated. An equal volume of molten 1% SeaPlaque GTG agarose (Lonza) at 42°C was added to the worm solution and approximately 80μl of the solution was pipetted into agarose plug molds (Bio-Rad #1703713), on ice. Plugs were allowed to solidify at 4°C for at least 10 minutes. Plugs were extracted to 2mL tubes and worms were anesthetized in 300μL TEL [9mM Tris, 90mM EDTA pH 8, 10mM levamisole] on ice for 30 minutes. TEL was replaced with 300μL lysis buffer [1% SDS, 1mg/mL Proteinase K (Sigma-Aldrich P4850), 8mM Tris, 80mM EDTA pH 8, 1mM levamisole] and worms were digested for 24 hours at 50°C. After digest, plugs were transferred to 24-well plates, lysis buffer was removed, and plugs were rinsed in 300μL TE [10mM Tris, 1mM EDTA]. Plugs were then washed with TE at least eight times for 20 minutes each at room temperature, with at least one wash being performed overnight at 4°C. Plugs were then stored at 4°C in TE until use.

Plugs were prepared for CHEF gel electrophoresis as follows: plugs were equilibrated in 1X NEB 3.1 buffer by soaking either overnight at 4°C or on the day of digestion for ≥1 hour in 1X NEB 3.1 buffer then replacing with fresh 1X NEB 3.1 buffer in a 24-well plate on ice. Approximately ¼ of the plug was cut with a razor blade and transferred to a parafilm-wrapped slide. In-plug restriction digest was used to cleave the rDNA array from the rest of chromosome I; SwaI cuts 3927bp upstream of *rrn-3.56* and not at all within the rDNA. 4µl SwaI was added to the surface of each plug fragment. Plugs were placed in a small humid chamber in a 25°C incubator and incubated for 4 hours for digest to occur before loading into the CHEF gel. CHEF gels were prepared by placing digested plugs and agarose-embedded ladders onto the teeth of gel combs. The following ladders were used: Yeast Chromosome PFG Marker (*S. cerevisiae*) (NEB #N0345; discontinued); *Hansenula wingei* chromosomes (maximum size 3.13 Mb, Bio-Rad 170-3667). 0.8% agarose prepared in 0.5X TBE at 55°C was poured around the plugs and solidified for at least 30 minutes. The gel was transferred to a CHEF gel box (Bio-Rad CHEF DRII), and run with the following conditions: for long rDNA arrays (total size ∼1Mb-3.13Mb): 100V for 68hr, 14°C, switch times = 300-900s. For medium-length rDNA (total size ∼225kb-1.1Mb): 165V for 66 hours, 14°C, switch times = 47-170s. After run completion, gel was stained by soaking in ethidium bromide (0.3ug/mL in 0.5X TBE) to visualize the ladders and imaged on a Bio-Rad GelDoc XR+. Southern blotting was performed as described previously (22,120). Briefly, each gel was washed 2x for 10 minutes in 0.25N HCl to nick and depurinate DNA, washed 2X for 15min in 0.5N NaOH, 1M NaCl, then washed 1X 30min in 0.5M Tris, 3M NaCl. DNA was then transferred from the gel to a nylon membrane overnight (Perkin Elmer GeneScreen Hybridization Transfer Membrane) and crosslinked to the membrane using a Stratagene Stratalinker UV Crosslinker. The probe for Southern blotting covers an 850bp region that overlaps the first 300bp of *rrn-1* (18S) amplified with primer pair EM50+EM51 (**Table S7**) and purified with the Zymo Clean & Concentrator kit (D4013) before radioactive labeling (22).

To determine rDNA array size, the distance between the bottom of the well and the middle of a sample band was measured with a ruler for the presented Southern blots. The distance between the bottom of the well and each of the ladder bands was measured from an image of the ethidium bromide stained gel taken before membrane transfer. The relationship between band size and gel distance was plotted in Microsoft Excel and calculated band sizes were divided by 7.2kb (the size of a single rDNA repeat unit) to determine rDNA copy number (**File S10**).

### Worm assays

For all assays, prior to assay initiation, worms were maintained on NGM + OP50 plates unless otherwise specified. All assays were performed on worms that were at least three generations removed from starvation, freezing, or contamination, unless otherwise specified.

#### Competitive fitness assays

Ten L4 worms of each of the two genotypes to be competed were picked to a 15cm NGM high peptone plate (20g/L peptone) seeded with NA22. Three to five 15cm plates were set up per competition trial, and three to four trials were conducted per genotype pair. Trials for some genotype pairs were conducted simultaneously (where indicated in figure legends), meaning that 4-5 competition plates of one set of genotypes were set up and propagated on the same days and using the same media batches as 4-5 competition plates for another pair of genotypes (ex. four plates of N2 competed against SEA51 alongside four plates of SEA300 competed against SEA51). Within a trial replicate plates were maintained as separate independent propagations. The initial plates were allowed to grow to starvation (∼6-7 days), after which a chunk of agar approximately 3cm x 4cm was cut out and propagated onto a new NGM high peptone + NA22 plate. This was done for seven plates subsequent to the first plate, with plates reaching starvation between each propagation (note: infrequently plates were propagated when only near starvation, if the other plates in the trial had starved). A final plate propagation was done through liquid transfer: starved worms from the eighth plate were washed off in 1X M9, spun 1500rpm 1 min, washed once in 1X M9, spun 1500rpm 1 min, and brought down to a volume of ∼3mL. L1 density was determined and ∼12,000 L1 were plated on a final NGM high peptone + NA22 plate, which was analyzed when these worms reached ∼L4 stage. We approximate the initial plate to have starved after 2-3 generations, and each subsequent plate after ∼1 generation, thus equaling ∼10-11 generations of competition.

Each competition included the test strain SEA51 (*mIs13[myo-2p::GFP + pes-10p::GFP + F22B7.9p::GFP]*) to allow for tracking via transgene GFP fluorescence. We quantified population proportions of GFP positive and negative worms using a COPAS Biosort (Union Biometrica). Two days after L1s were seeded on the final plate, worms were washed off with 1X M9, spun 1500rpm 1 min, washed once in 1X M9, spun 1500rpm 1 min, and brought up to a ∼10mL volume with 1X M9. Worms suspended in 1X M9 were quantified for size and fluorescence by flowing through the COPAS, collecting data for at least 1,000 worms per plate. Objects were gated for size (time of flight) to enrich for L4/adult. Thresholds of green fluorescence peak height were used to call GFP presence or absence: a green fluorescence peak height greater than 50000 indicated GFP presence, a green fluorescence peak height of less than 20000 indicated GFP absence. Very few worms fell in the expanse between these two values (0 worms for most samples, maximum 3 worms per sample, see **File S14**), indicating confidence in GFP status calling. Controls of the parental strains were also quantified by COPAS to confirm accurate GFP detection. Most paired competitions also included COPAS quantification at an early time point: after the competition had been chunked to plate 2 to continue propagation, the starting plate was also washed off with 1X M9 as described above and ∼12,000 L1 worms were plated on a 15cm high peptone plate. This early population was allowed to grow for 2 days at 20°C before being quantified for population proportions with the COPAS Biosort. Proportions at this timepoint were on average 54% with a standard deviation of 8% (**Figure S5**).

#### Early life fertility

Worms were synchronized by allowing 10 Day 1 adult worms to pulse-lay for 1 hour on NGM + OP50. Two days later, single worms were picked to individual 3cm plates. For some assays, 20 individual worms were singled to 3cm plates and all plates were scored (**Figure S6B**). For others, 30 individual worms were singled to 3cm plates and 20 plates per strain were randomly selected to be scored (**Figure 3C, Figure S6A**). The latter measure was implemented to reduce bias in the exact stage of worm selected, as some variation in larval size was present on each pulse-lay plate, despite the short pulse-lay duration. Either 90 hours (**Figure S6B**) or 86 hours (**Figure 3C, Figure S6A**) after initiation of the pulse-lay, adults were removed from the 3cm plates. Progeny were allowed to grow for 48-72 hours before counting number of viable progeny (**File S15**). Plates were stored at 4°C when counting over multiple days, to prevent interference from the next generation of worms.

#### Cuticle permeability

Cuticle permeability assay was based on two previous publications (121,122). Briefly, approximately 30 Day 1 adult worms were picked to a watchglass slide containing 150μl 1X M9 with 100mM levamisole and 10μg/mL Hoechst 33258 dye (Sigma) to simultaneously stain and anesthetize the worms. Worms were incubated in the stain for 30 minutes statically in the dark, then washed 5x with 100μl 1X M9 in the watchglass slide. Worms were picked into a spot of ∼10μl 1X M9 on a 2% agarose pad on a microscope slide with an eyelash pick, recovering approximately 20-25 worms per strain. Hoechst stain was visualized with the DAPI filter on a Zeiss Apotome, and the number of worms with nuclear staining of hypodermal cells was quantified.

#### Lifespan

Lifespan assays were conducted with worms propagated at least three generations away from starvation, freezing, or contamination. Ten L4 worms per strain were picked to 3cm plates seeded with OP50, for a total of 50 worms per strain, unless otherwise noted. Within a given assay, strain identities were anonymized at the beginning of the experiment so that personnel performing the scoring were blind to genotype for the length of the assay. Starting on Day 2 of adulthood, worms were transferred to new 3cm plates daily until egg-laying ceased. Viability was determined by visible movement of the worm, either spontaneously or in response to plate tapping or gentle touch with a wire pick. Lack of response to repeated prodding by a wire pick declared as dead (**File S16**). Analysis of lifespan data was performed in R (package survival).

#### Male frequency assay

Worm strains with various rDNA copy numbers all carrying the *him-5(ok1986)* allele were synchronized by pulse-lay of Day 1 adults. Two days later, five healthy L4 individuals per strain were singled to 3cm NGM plates seeded with OP50. Adult worms were transferred to new 3cm NGM + OP50 plates every 24 hours for 3 days (3 total transfers). Twenty-four hours after the adult was removed, each plate was scored for the presence of unhatched (dead) embryos. Forty-eight to seventy-two hours after the adult was removed, hatched progeny were counted and scored as: healthy hermaphrodite, male, developmentally delayed (too young to discriminate between males and hermaphrodites), or dumpy hermaphrodites. Males, developmentally delayed worms, dumpy hermaphrodites, and dead embryos were categorized as evidence of nondisjunction events (**File S17**).

### RNA-sequencing

#### Preparation of worms

Large pools of *C. elegans* were grown by bleaching an asynchronous worm population and allowing embryos to hatch and starve out for 20-24 hours statically on 10cm unseeded NGM plates overlaid with 10mL 1X M9. Starved L1 worms were transferred to 15mL conical tubes and washed once in 10mL 1X M9. 15,000 worms per strain were plated on individual 15cm high peptone NGM + NA22 plates and grown to the first day of adulthood (∼72 hours post-plating). Worms were harvested into 15mL conical tubes by washing the 15cm plates with 15mL 1X M9. Worms were allowed to settle for 3 minutes, supernatant was removed, and worms were washed 4x in 12mL 1X M9. 500μl Trizol was added to the worm pellet, the mixture was transferred to a 1.5mL tube, flash frozen in liquid nitrogen, and stored at −80°C.

#### RNA isolation

RNA was isolated with a Trizol-chloroform extraction protocol. The stored worm/Trizol mixture was freeze-thawed 3x between 37°C and a liquid nitrogen bath. 200μl additional Trizol was added to the worm/Trizol mixture and mixed by pipetting up and down, then incubated for 5 minutes at room temperature. 140μl chloroform was added, and samples were mixed by shaking vigorously for 20 seconds. The mixture was incubated for 2 minutes at room temperature, then spun 15 minutes at 12,000xg at 4°C. The supernatant was transferred to a clean microcentrifuge tube containing 140μl chloroform; tubes were mixed by shaking vigorously for 20 seconds and allowed to sit for 2 minutes at room temperature, then spun 15 minutes at 12,000xg at 4°C. Supernatant was transferred to a Lo-bind tube containing 300μl of isopropanol and tubes were inverted gently to mix. Samples were incubated 10 minutes at room temperature, then spun 10 minutes at 12,000xg at 4°C. Supernatant was removed, and pellet washed with 600μl cold 75% ethanol. Pellets were allowed to dry 5-10 minutes, then resuspended in 38μl Invitrogen RNase-free water and resuspended at room temperature.

The 38μl RNA was treated with TURBO DNase, adding 2μl TURBO DNase, 5μl 10X TURBO buffer, and 5μl 100mM MnCl_2_. DNase treatment was performed for 1 hour at 37°C. RNA was then purified with an ethanol precipitation, and the final RNA pellet was resuspended in 100μl Invitrogen RNase-free water. RNA quality was assessed with TapeStation (Agilent); RNA concentration was determined with nanodrop. All RNA samples were stored at −80°C.

### Library preparation

The library preparation protocol was derived from a previously-published bulk sci-RNA-seq protocol with an added RNase H-based rRNA depletion step (123–125). For rRNA depletion, 1μg RNA was combined with 1μg rRNA-complementary oligoPool (IDT, 50nt non-overlapping fragments complementary to processed rRNA transcripts, covering length of each rRNA) and brought to a volume of 5μl with 1X hybridization buffer (200 mM NaCl, 100 mM Tris pH7.4). The mixture was incubated at 95°C for 2 minutes, then brought to 45°C with a ramp speed of −0.1°C/s. 5μl RNase H mixture (5U Hybridase Thermostable RNase H (Epicentre), 0.05μmole Tris HCl pH 7.5, 1μmole NaCl, and 0.2μmole MgCl_2_) preheated to 45°C was added to the hybridized RNA/oligo mixture and mixed by pipetting. Digestion was performed for 45 minutes at 45°C.

rRNA-depleted RNA was purified with a 2.2X RNAClean SPRI bead treatment (Beckman Coulter). Purified, rRNA-depleted RNA was treated with Turbo DNase (Invitrogen) to remove the rRNA-hybridizing DNA oligos: 0.1 volumes of 10X TURBO DNase Buffer was added to the RNA, followed by 2μl TURBO DNase. The mixture was incubated at 37°C for 45 minutes. DNase-treated RNA was cleaned with 2.2X RNA SPRI beads. Final elution volume was 12μl, of which 9μl was recovered and used directly in the next step (oligo-dT incubation).

To the 9μl of final rRNA-depleted RNA, 1μl 10mM dNTPs and 2μl 25μM oligo-dT(VN) (IDT, containing UMIs) were added and the mixture was incubated 5 minutes at 65°C. Reverse transcription was performed by adding 4μl SuperScript IV Buffer, 2μl 0.1M DTT, 1μl SuperScript IV Reverse Transcriptase, and 1μl SUPERase-IN RNase inhibitor and incubating at 42°C for 50 minutes then 70°C for 15 minutes. Second strand synthesis was performed with the NEBNext Ultra II Non-Directional RNA Second Strand Synthesis Kit. 2μl reverse transcription product was combined with 6.5μl Invitrogen RNase Free water, 1μl NEB Second Strand Synthesis Buffer, and 0.25μl NEB Second Strand Synthesis Enzyme Cocktail and incubated at 16°C for 150 minutes then 75°C for 20 minutes. The resulting cDNA was then either used immediately or stored at 4°C overnight for use the next day.

Libraries were sequenced using a Nextseq 550 with a 75 Hi kit (Illumina). Read lengths used were: Index 1: 10bp, Index 2: 10bp, Read 1: 18bp, Read 2: 52bp.

#### Data analysis

Reads were demultiplexed with bcl2fastq (version 2.20). UMI information from Read 1 was merged into the Read 2 file for use in later steps. Poly-A tails were trimmed with Trim Galore (version 0.6.6), with accessory modules perl (version 5.24.0), python (version 2.7.13) and cutadapt (version 1.18). Trimmed reads were aligned to the WS260 *C. elegans* reference genome using STAR version 2.6.1c. Reads were filtered for ambiguously-mapping reads and sorted with samtools (version 1.9). PCR duplicate reads were identified with the UMI from the oligo-dT and removed with a custom script (124). Each of the previous steps was performed independently for each sequencing run performed. To merge reads from multiple sequencing runs, Samtools was used to merge duplicate-removed bam files. Duplicate removal was then repeated. The number of reads aligning to the rRNA was counted with bedtools (version 2.29.2).

A file of counts per gene region of interest was produced with HTSeq (version 0.12.4) and accessory modules python (version 3.7.7), numpy (version 1.19.2), samtools (version 1.10), and pysam (version 0.16.0.1). The commands htseq-count -m union -i gene_id -r pos -a 10 -- stranded=yes were used, with the WS260 canonical geneset gtf file used to provide gene coordinates, and final counts were merged and output into a counts file. The counts file was imported into R version 4.0.4 for analysis of differential gene expression with DESeq2. Gene Set Enrichment Analysis of RNA-seq data was performed with the WormBase tool with a q value threshold of 0.1 (126) (**Files S18-S21**).

### rRNA quantification

#### TapeStation

The same sample of RNA that was extracted for the RNA-seq analysis described above was used for the 18S and 28S rRNA quantification by TapeStation capillary electrophoresis. 100ng RNA was used for each sample and the standard Agilent RNA ScreenTape protocol was used (**File S11**). Total RNA was normalized before loading, and the TapeStation was used to measure the integrated area of the bands corresponding to each rRNA species.

#### RNA gel

The same sample of RNA that was extracted for the RNA-seq analysis described above was used for the RNA gel. RNA was diluted to 500ng/μl and 5.5μl RNA was denatured at 65°C for 5 minutes with 10μl 96% formamide in a 20μl reaction containing 3.5μl 6X DNA loading dye and 1μl 10mg/ml ethidium bromide. After denaturing, the sample was immediately put on ice for 5 minutes, then 18μl of the sample was loaded into a 1.2% agarose gel prepared in fresh 1X TAE buffer. Gel electrophoresis was performed for 7 hours at 80V, and the resulting gel was imaged on a ChemiDoc MP (Bio-Rad). Band intensities were quantified in ImageJ (**File S12**).

#### RT-qPCR

200ng RNA was used for reverse transcription in a 20μl reaction with RevertAid Reverse Transcriptase kit (Thermo Scientific). Reverse transcription was performed at 25°C for 5 min, 42°C for 60 min, and 70°C for 5 min, and the samples were stored at −80°C until use in qPCR reactions. Because rRNA is highly abundant, the reverse transcription product was diluted for the qPCR reactions to an equivalent RNA amount of 0.2ng/μl, and 5μl of this 0.2ng/μl RNA equivalent was used in each qPCR reaction. For each set of primers used in each qPCR plate, a standard curve was set up, diluting the original RT product 1:5 a total of six times. For the standard curve, 5μl of each dilution was used in the qPCR reactions. The qPCR reactions were performed with Roche LightCycler 480 SYBR Green I Master Mix in 20μl reactions, with 5μl 10μM of each primer per reaction. qPCR reactions were performed in a BioRad CFX Connect with the following conditions: 95°C 10 min, *95°C 10s, 60°C 15s, 72°C 30s, image, repeat from * a total of 40x. Results were normalized to actin (127). Primers used are listed in **Table S7**. qPCR analysis was performed in Microsoft Excel and plotted in R with ggplot2 (**Files S13**).

### Statistics and data visualization

All statistical tests described (including ANOVA, Tukey’s test, Shapiro-Wilk normality test, Wilcoxon test, and t-tests, as indicated in figure legends) were performed in R (128). Data were visualized in R with the base plotting system or with ggplot2 (128,129). The Color Universal Design palette (130) was used for some visualizations. R libraries used include R/DESeq2, dplyr, extrafont, ggplot2, ggpubr, ggsignif, plotrix, qtl, rstatix, splines, survival, and tidyverse (129,131–137).

## DATA AVAILABILITY

Strains (**Table 1**, **File S1**) are available upon request. VCF files are available in supplemental information (**File S3-S9**). FASTQ files for sequenced RILs, NILs and RNA-sequencing are available through SRA, BioProject PRJNA1182379.

## ACKNOWLEDGEMENTS

We would like to thank the Moerman lab and Teotonio lab for kindly providing wild worm isolates. Some strains were provided by the CGC, which is funded by NIH Office of Research Infrastructure Programs (P40 OD010440). We would like to acknowledge members of the Queitsch, Waterston, and Brewer/Raghuraman for helpful discussion. We thank the Kaeberlein lab for use of their COPAS Biosort. This work was supported by funding from NIGMS (grants 5R01GM122088 and R35 GM139532 to C.Q.), NHGRI (grant 1RM1HG010461 to C.Q), NSF (award 2437133 to E.A.M.), and NIA (F31 AG063450 to A.N.H.).

## FIGURE LEGENDS

**Figure S1: Haplotype blocks of RILs.**

**A:** Top: The genotypes of the 58 RILs with ∼130 rDNA copies are presented, with yellow representing haploblocks matching the parental MY1 wild isolate genotype and green representing the parental SEA51 N2-derivative genotype. Bottom: The genotypes of the 60 RILs with ∼417 rDNA copies are presented. The black carat indicates the rDNA locus at the end of chromosome I. Genotypes were determined with GATK HaplotypeCaller and the map was filled with the max marginal method. **B:** Short-read sequencing-based 45S rDNA copy number estimates are given for RILs pre- and post-propagation for 20 generations. Data are the same as presented in Figure 1C and D but plotted here on a continuous axis. Pink lines indicate parental control strains (see Figure 1).

**Figure S2: CHEF gel validation of rDNA copy number.**

**A:** Six RILs with rDNA linked to the *mIs13* transgene were analyzed by CHEF gel and Southern blot under conditions to resolve rDNA arrays near 100 copies in length (see **Table S1**, **Methods**). Band sizes were calculated based on distance of band migration from the well compared to an *S. cerevisiae* chromosome ladder that was visualized by ethidium bromide staining prior to Southern blotting (**File S10**). Copy numbers were calculated from the base pair size of the band divided by 7.2kb, the size of a single *C. elegans* rDNA unit. **B:** N2 and NIL rDNA arrays were separated with conditions that resolve arrays near 100 rDNA copies. Copy numbers were calculated as in A. N2 is in lane 1 and allele identifiers for introgressed rDNA arrays in NILs are indicated in subsequent lanes. **C:** NILs with high rDNA copy number were analyzed with CHEF gel conditions that resolve arrays larger than 200 copies. Copy numbers were calculated based on reference to an *H. wingei* chromosome ladder that was visualized by ethidium bromide staining prior to Southern blotting. Allele identifiers for rDNA arrays are indicated. Numbers on the blots indicate the calculated rDNA copy number for each band.

**Figure S3: 45S rRNA levels in NILs do not differ in a copy-number-dependent manner.**

**A:** Equal quantities of RNA extracted from Day 1 adult worms were denatured and run on an agarose gel for each sample. Each lane is a separate biological replicate. **B** and **C:** Intensities of 28S and 18S bands from **A** were quantified with ImageJ. **: p<0.01 as determined by ANOVA and Tukey’s HSD. **D-F:** Steady-state rRNA levels in NILs. **D**: 18S rRNA levels in NILs were measured by TapeStation. No significant differences in 18S rRNA levels were observed between any strains as assessed by ANOVA and Tukey’s HSD. **E**: 28S rRNA levels in NILs were measured by TapeStation. Data are not normally distributed as determined by the Shapiro-Wilk normality test. No significant differences in 28S rRNA levels were observed between any strains as measured by pairwise Wilcoxon test and Benjamini-Hochberg significance adjustment. **F**: 5S rRNA levels in NILs were measured by RT-qPCR, normalized to actin. No significant differences in 5S rRNA levels were observed between any strains as measured by ANOVA and Tukey’s HSD. Legend at the right indicates the rDNA allele for each strain in D-F. **G:** Diagram of 45S pre-rRNA processing in *C. elegans*, adapted based on Wu *et al*. 2018 (138). Primer pairs are as indicated in **Table S7**. **H:** Pre-rRNA levels were quantified by RT-qPCR in NILs with large regions of linked wild isolate DNA (see Figure 2A). rRNA levels are normalized to actin. Error bars are mean ± standard deviation. No significant differences are present between any NILs and N2 (ANOVA with Tukey Honest Significant Difference test).

**Figure S4: The 420-rDNA strain with large wild isolated linked DNA (allele *catIR16*) exhibits a competitive fitness defect that is not as severe as that of the 417-rDNA strain.**

Competitions of strains with high rDNA copy number (GFP(-); alleles indicated at bottom) set against SEA51 (GFP(+); 130 rDNA copies). For panels A-C, each bar shows the relative proportion of worms that are GFP(+) (green) or GFP(-) (white) after ∼10-11 generations of competition. At least 1,000 worms were quantified to determine the proportion of each bar (**File S14**). **A:** Five independent replicates were conducted of SEA51 competed against N2, performed at the same time as four replicates of SEA51 competed against the 420-rDNA NIL (allele *catIR16*, which has ∼3.3Mb of wild isolate DNA linked to the rDNA array (Figure 2A)). **B:** Experiment set up similarly to A. Five replicate competitions between SEA51 and N2 were performed, with four replicate competitions between SEA51 and 420-rDNA NIL (allele *catIR16*) performed at the same time. **C:** Competitions between SEA51 and each of the two strains with high rDNA copy number were performed side-by-side, with four independent replicates each. High rDNA copy number strains are the 417-rDNA NIL (allele *catIR12*) and 420-rDNA NIL (allele *catIR16*). **D:** A simulation of selection coefficients was used to determine what strength of selection is required to give a certain population proportion after 11 generations of propagation with a propagated population size of 1000.

**Figure S5: Early in the competitions, strains are at near-equal proportions.**

Competition population proportion data was collected early in the assay, corresponding to the point when the populations were transferred for the first time (out of eight total transfers in the assay), approximately 2-3 generations into the assay. **A:** Early timepoint data for the competitions presented in Figure 3B. Four plate replicates each are shown for SEA51 competing against either N2, the 417-rDNA NIL (allele *catIR12*), or the 420-rDNA NIL (allele *catIR29*). These three sets of competing pairs were assayed simultaneously. **B:** Early timepoint data for the competitions presented in **Figure S4A**. Data are shown for SEA51 competing against either N2 (five replicates) or the 420-rDNA NIL (allele *catIR16*) (four replicates), propagated at the same time. **C:** Early timepoint data for the competitions presented in **Figure S4B**. Data are shown for SEA51 competing against either N2 (five replicates) or the 420-rDNA NIL (allele *catIR16*) (four replicates), propagated at the same time. **D:** Early timepoint data for the competitions presented in **Figure S4C**. Competitions against SEA51 are shown for two strains with high rDNA copy number (417-rDNA NIL (allele *catIR12*) and 420-rDNA NIL (allele *catIR16*)), propagated at the same time. For all panels, each bar shows the relative proportion of worms that are GFP(+) (green) or GFP(-) (white) after ∼10-11 generations of growth. At least 1,000 worms were quantified to determine the proportion of each bar (**File S14**).

**Figure S6: Fertility phenotyping of N2 NILs reveals no copy-number-dependent differences.**

**A:** Early life fertility of NILs was compared to N2 in three replicates, n=20 individual worms per strain per replicate. These strains represent the panel of NILs with minimal rDNA-linked wild isolate DNA. The data fail the Shapiro-Wilk normality test and are not normally distributed. A nonparametric Scheirer Ray Hare test of Progeny by Strain and Replicate shows a significant effect of Strain (p=0.00137) and Replicate (p<0.00001) across the three replicates. Due to the lack of normality and the significant effect of replicate, strain-by-strain comparisons were performed separately for each replicate. The data for N2, *catIR12*, and *catIR29* are the same as the data presented in Figure 3C. **B:** Two replicates of the early life fertility assay were performed on the panel of NILs with large regions of rDNA-linked wild isolate DNA, n=20 individual worms per strain per replicate. The data fail the Shapiro-Wilk normality test and are not normally distributed. For both panels **A** and **B**, statistical tests represented in the figure are Pairwise Wilcoxon tests with Benjamini-Hochberg procedure performed to compare strains. * p<0.1, ** p<0.05, *** p<0.01, **** p< 0.001.

**Figure S7: Lifespan phenotyping of NILs reveals no reproducible copy-number-dependent differences.**

**A** and **B**: Lifespans of NILs with large linked wild isolate DNA regions (see Figure 2) were compared to N2. Lifespans for all five strains in **A** and **B** were performed simultaneously (panels **A** and **B** present the same N2 data)(n=50). **C** and **D**: Lifespans of NILs with minimal linked wild isolate DNA (see Figure 2) were compared to N2 in biological replicates of the experiment presented in Figure 4. **C**: (n=50). **D**: (n=40 (N2), n=44 (rDNA-417), n=42 (rDNA-420)). A significant difference (p = 0.01) was detected between N2 and rDNA-73 in the replicate presented in panel **C** only.

**Figure S8: Visible signs of nondisjunction events do not differ with different rDNA copy numbers in NILs carrying the *him-5*(*ok1986*) allele.**

**A:** A mutation in a gene involved in meiotic break positioning, *him-5(ok1986)*, was crossed into the rDNA copy number NILs to create strains sensitized for nondisjunction events. Progeny were then scored for incidence of males, dumpy worms, dead embryos, and other defects such as slow growth. The percent of progeny exhibiting such phenotypes is collectively plotted here as “nondisjunction”. Progeny from five adult worms were assessed per strain. No significant differences between any strains were observed by t-test and Bonferroni correction. **B:** The data from A are parsed out to present only the percent of progeny that failed to hatch (dead embryos) in NILs with *him-5*(*ok1986*). Dead embryos are assumed to arise from autosomal nondisjunction events. No significant differences were observed between any strains by t-test and Bonferroni correction. **C:** The data from A are parsed out to present only the percent male progeny (X-chromosome nondisjunction) in NILs with *him-5*(*ok1986*). The 73-rDNA NIL (allele *catIR28*) differed significantly from wild type in the percent male progeny, p<0.01. **D-F:** A second replicate was conducted as in A-C, assessing only N2 and the 73-rDNA NIL. No significant differences were observed between these strains in collective nondisjunction events (**D**) or the individual contributions of dead embryos (**E**) or male frequency (**F**) (student’s t-test).

**Figure S9: Cuticle permeability does not differ between N2 and the 73-rDNA NIL.**

Worms were stained with Hoechst, which does not normally penetrate intact worm cuticles. Worms that exhibited staining in their hypodermal nuclei were scored as having permeable cuticles. At least 13 worms were quantified for each replicate, three replicates per genotype (**File S22**). The *bus-8*(*e2698*) genotype was used as a positive control with known reduced cuticle integrity (139); in all three replicates of this genotype all worms had permeable cuticles. N2 and the 73-rDNA NIL (*catIR28*) have a fraction of worms with permeable cuticles and do not differ from one another in cuticle permeability, as measured by ANOVA and Tukey’s HSD. Error bars represent standard deviation of the percent permeable over the three replicates.

## Notes

### Competing Interest Statement

The authors have declared no competing interest.

### Summary of Updates

Additional analyses are included, additional figure panel in Fig. S1, updates to content.

