## Supplementary figures and images for "Phenotypic tolerance for rDNA copy number variation within the natural range of *C. elegans*"

### Supplemental Information

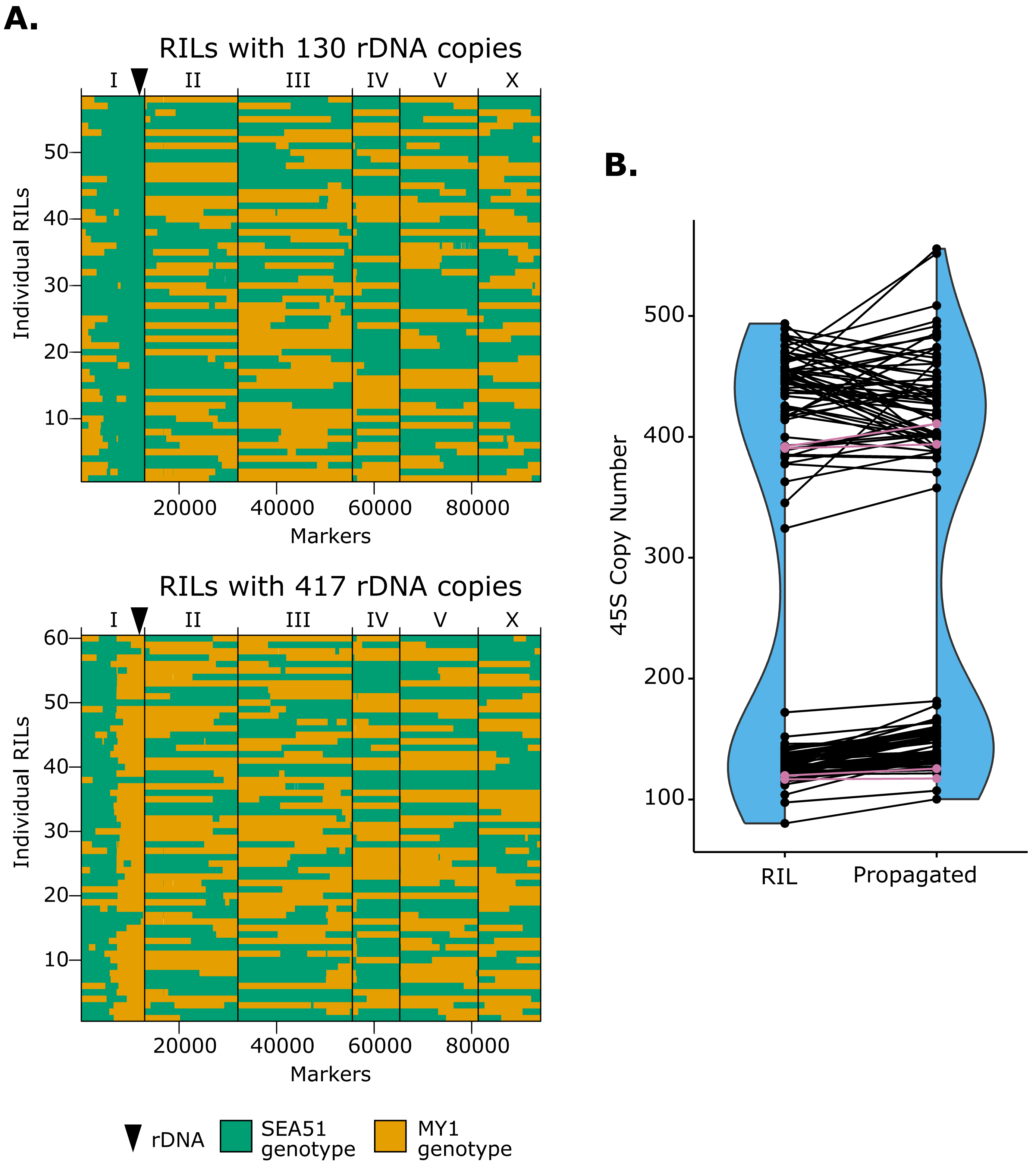

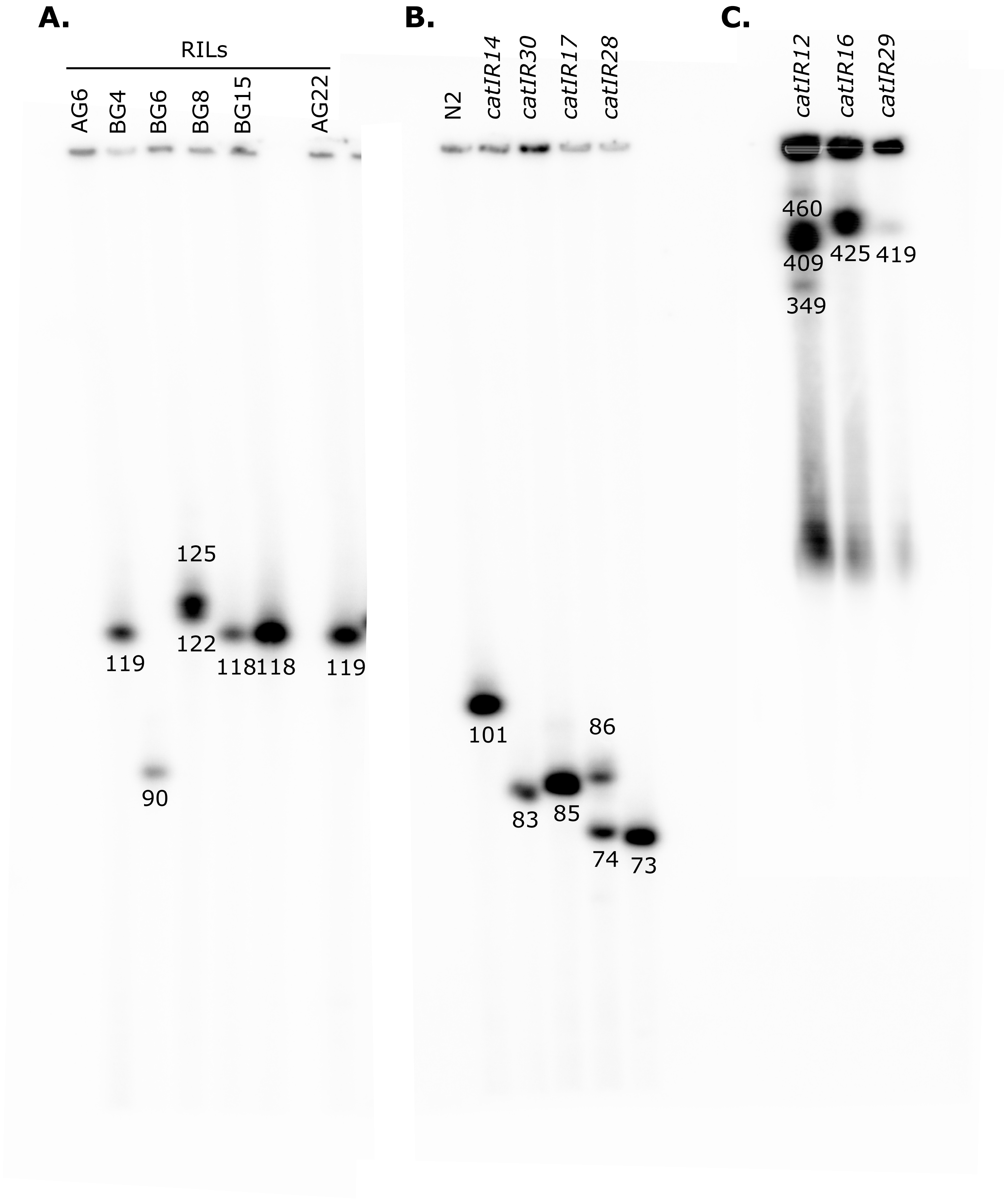

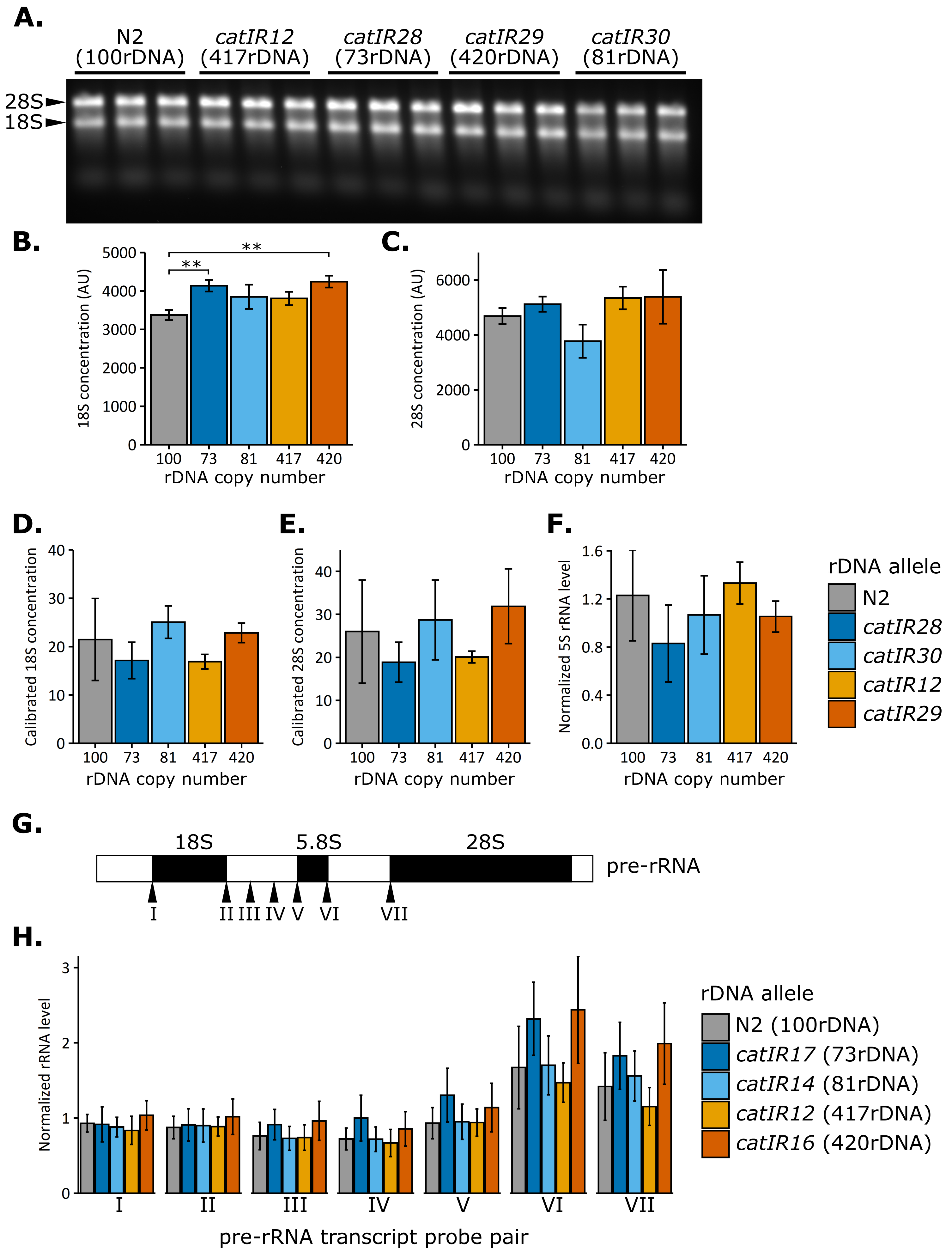

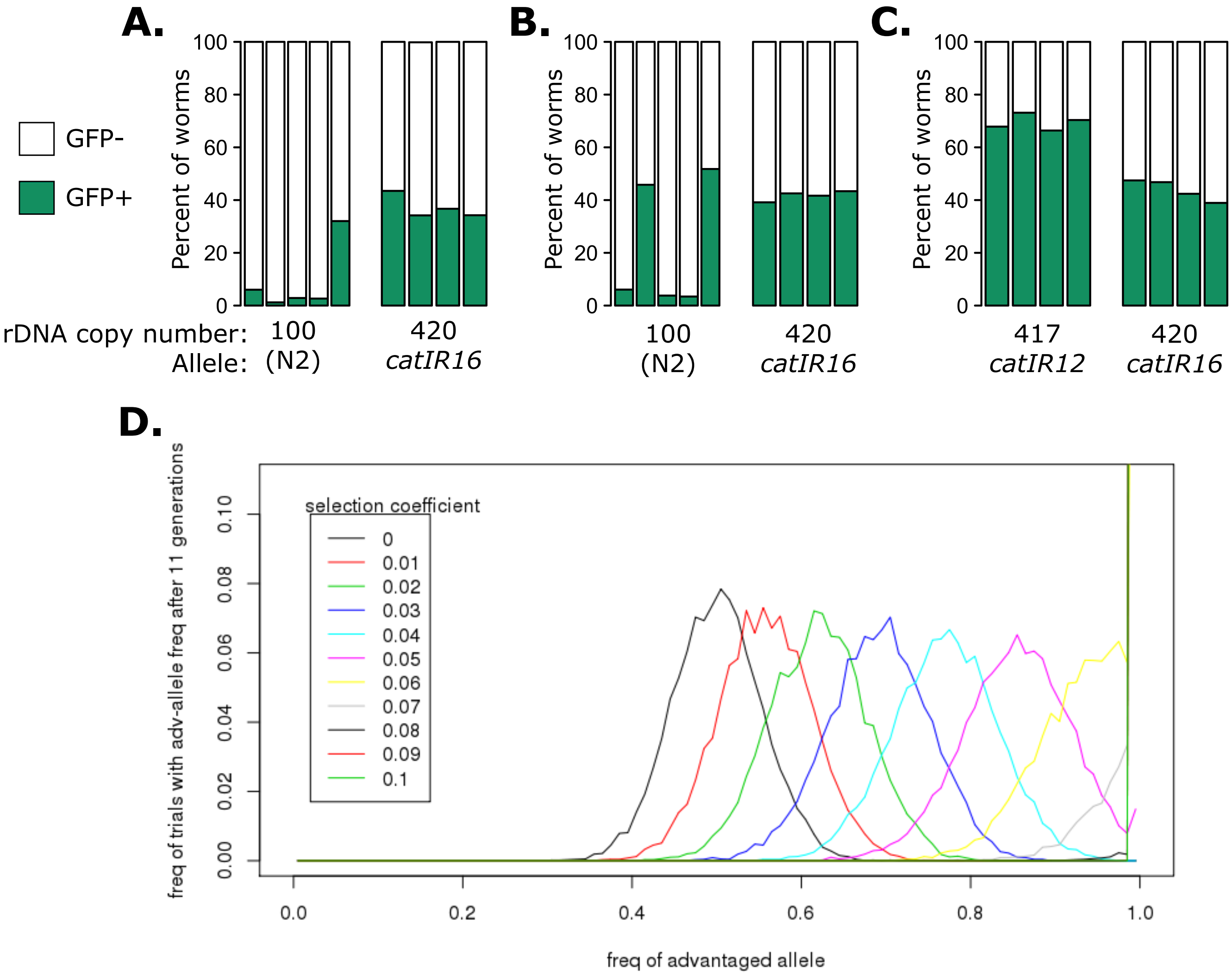

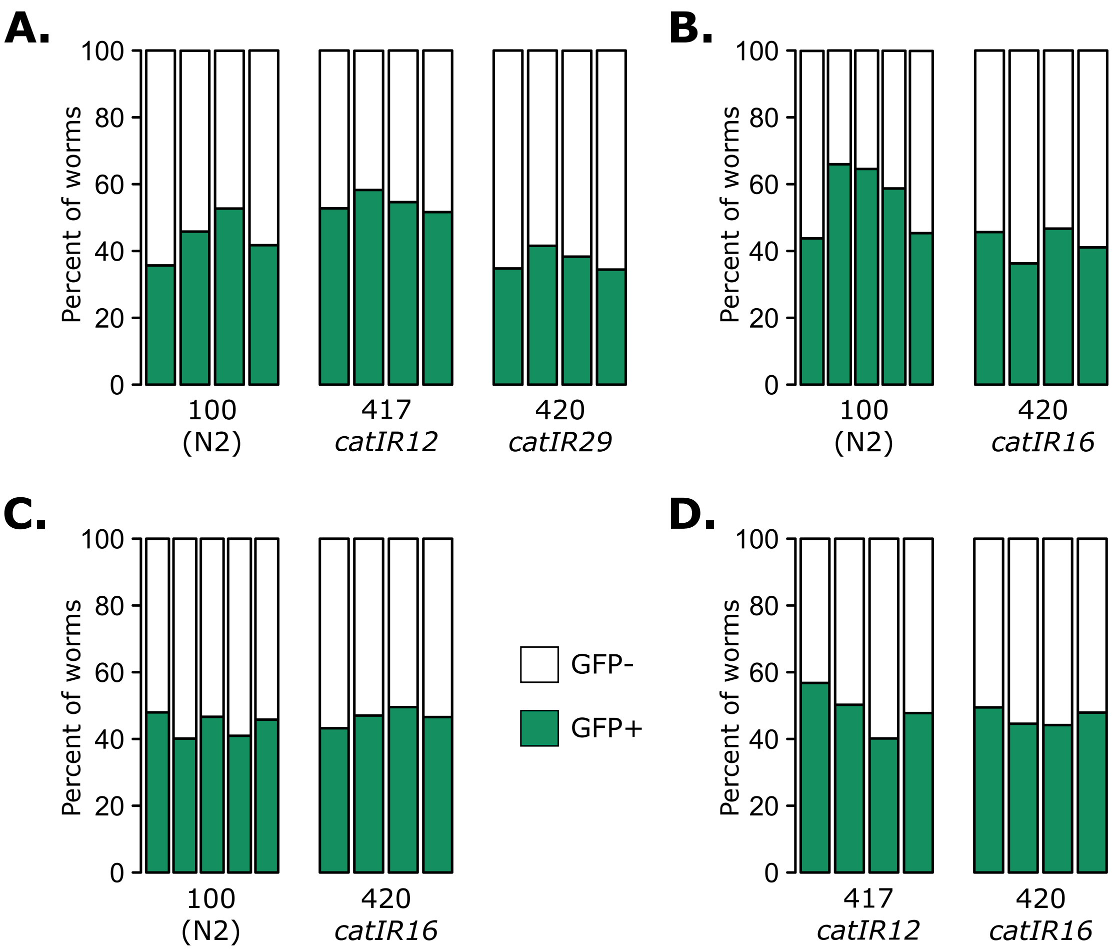

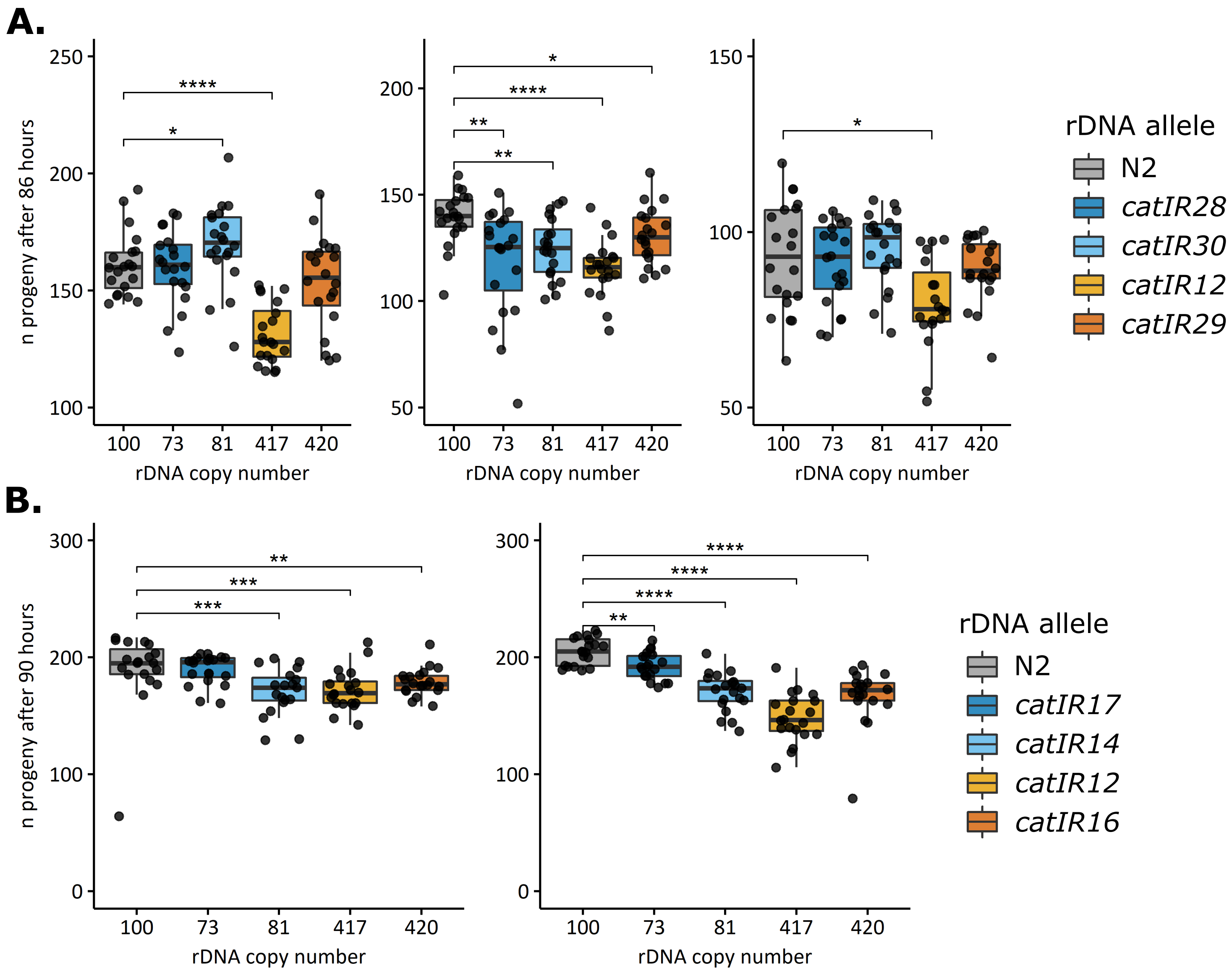

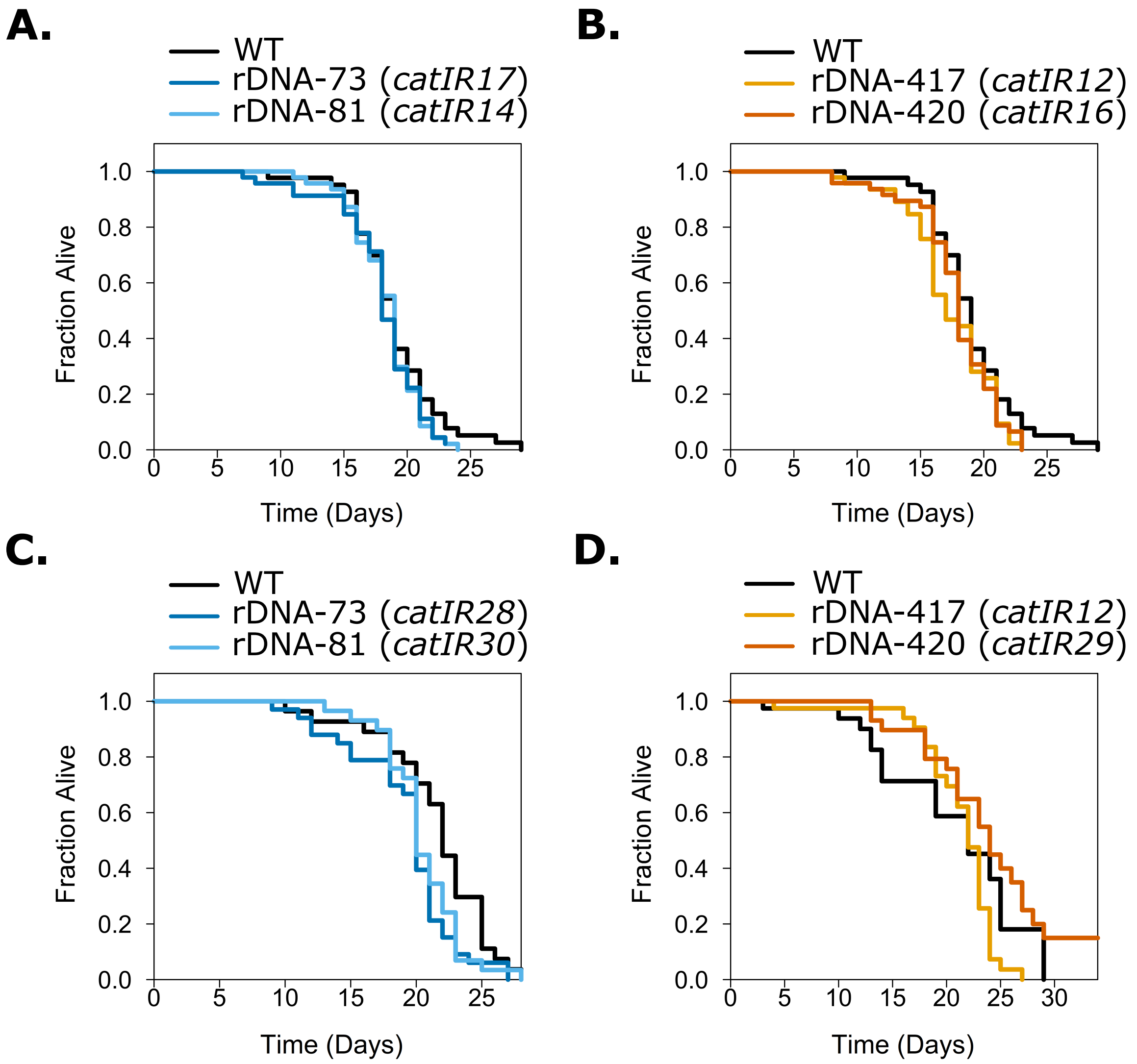

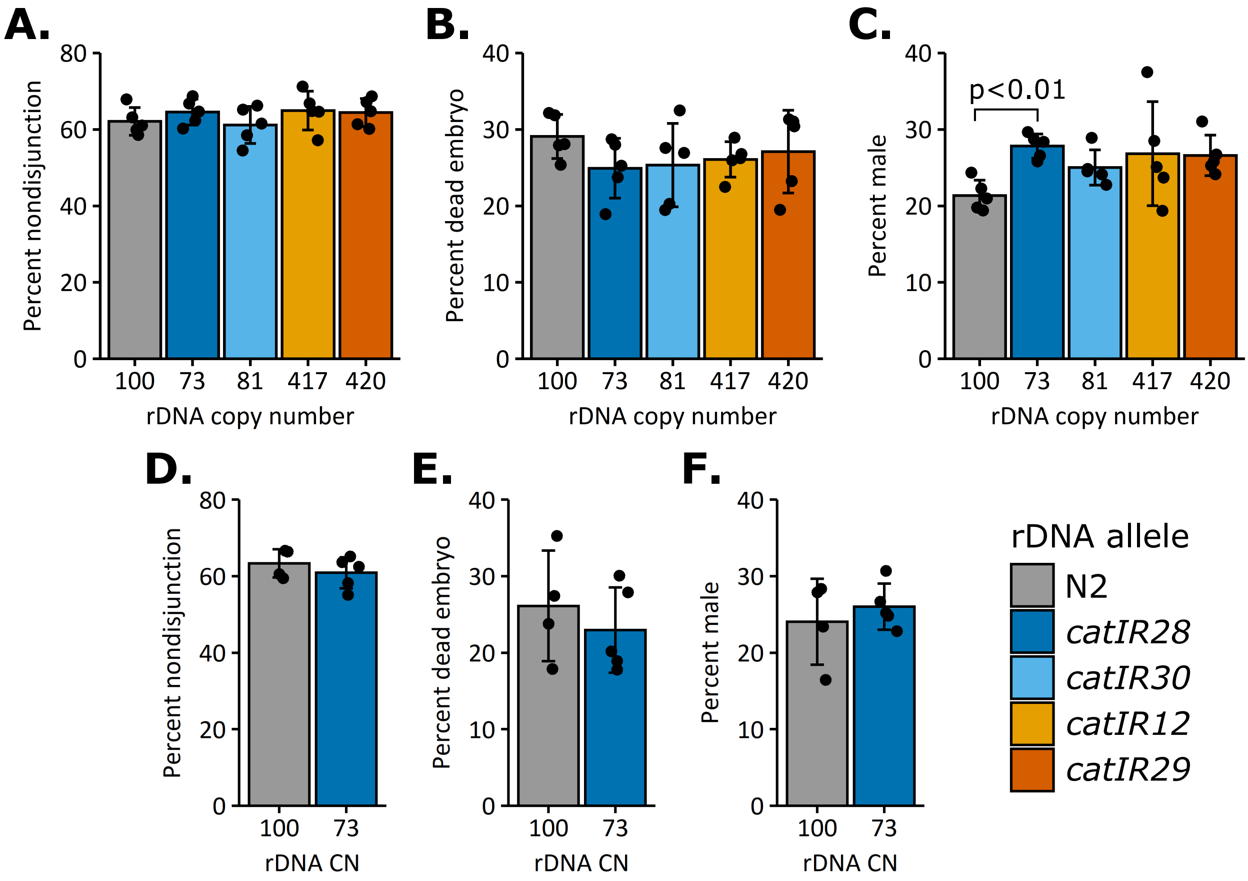

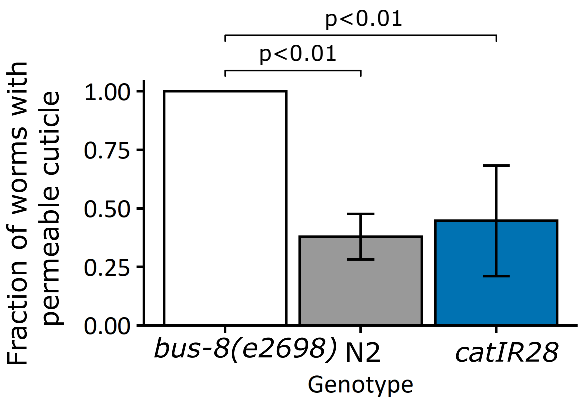
